# Tumor necrosis factor is a necroptosis-associated alarmin

**DOI:** 10.1101/2022.08.01.502280

**Authors:** Francesca Pinci, Moritz M. Gaidt, Christophe Jung, Dennis Nagl, Gunnar Kuut, Veit Hornung

## Abstract

Necroptosis is a form of regulated cell death that can occur downstream of several immune pathways. While previous studies have shown that dysregulated necroptosis leads to strong inflammatory responses, little is known about the identity of the endogenous molecules that trigger necroptosis-mediated inflammatory responses. Using a reductionist *in vitro* model, we found that soluble TNF is strongly released in the context of necroptosis. On the one hand, necroptosis promotes TNF translation by inhibiting negative regulatory mechanisms acting at the post-transcriptional level. On the other hand, necroptosis markedly enhances TNF release by activating ADAM proteases. In studying TNF release at single-cell resolution, we found that TNF release triggered by necroptosis is activated in a switch-like manner and exceeds steady-state TNF processing in magnitude and speed. Although this shedding response precedes massive membrane damage, it is closely associated with lytic cell death. In fact, we found that lytic cell death induction using a pore-forming toxin also triggers TNF shedding, indicating that the activation of ADAM proteases is not strictly related to the necroptotic pathway but associated with biophysical changes of the cell membrane upon lytic cell death. These results demonstrate that lytic cell death, particularly necroptosis, is a critical trigger for TNF release and thus qualify TNF as a necroptosis-associated alarmin.

## Introduction

Necroptosis is a lytic form of programmed cell death that can be triggered downstream of various immune receptors. Originally described to occur downstream of death receptors, necroptosis can be considered a “last resort type of response” of the cell, in that it is only engaged if other response pathways fail to be activated. This paradigm is well established in the context of TNF receptor 1 (TNFR1) signaling ^1^. Briefly, downstream of TNFR1 activation, Receptor-interacting serine/threonine-protein kinase 1 (RIPK1) can gain autophosphorylation activity when not ubiquitylated. This results in the recruitment of RIPK3 via homotypic RHIM domain interaction. RIPK3 then forms homo-oligomers, in which RIPK3 is phosphorylated, which in turn leads to the recruitment and phosphorylation of mixed lineage kinase domain-like pseudokinase (MLKL) ^2–5^. Upon phosphorylation, MLKL oligomerizes and translocates to the plasma membrane, where it mediates membrane permeabilization and rupture by a so far not fully understood mechanism ^6^. Besides RIPK1-dependent pathways, two additional RHIM domain-containing molecules can also trigger RIPK3 activation and thus necroptosis. Downstream of Toll-like receptors 3 and 4 (TLR3 and TLR4), RIPK3 can be activated by the adapter protein TRIF ^7,8^. In addition, the cytosolic sensor Z-DNA-binding protein 1 (ZBP1) can also recruit and activate RIPK3 ^9,10^. As for RIPK1, the engagement of RIPK3 is dependent on RHIM domain-mediated homotypic interactions ^11,12^. A well-known break on necroptosis induction is the catalytic activity of caspase-8 ^1,10,13^. In fact, caspase-8 plays an important role in cleaving and thus inhibiting RIPK1 and RIPK3 ^14,15^.

Being involved downstream of immune receptors, it is believed that necroptosis has evolved as a host defense mechanism against pathogens ^16^. In line with this notion, necroptosis is utilized as a “backup” cell death pathway when apoptosis is blocked by viral-encoded inhibitors ^2,16^. Moreover, virus-encoded inhibitors of the RHIM domain as well as decoy viral MLKL (vMLKL) that counteract the activity of key necroptotic mediators exist, confirming the importance of necroptosis in the control of infections ^17–19^.

Despite its beneficial contribution to pathogen clearance, necroptosis has also been reported to have a detrimental role in many inflammatory diseases under sterile conditions ^20,21^. The ability of necroptosis to promote inflammation is generally attributed to its lytic nature, which implies the release of its cellular content upon plasma membrane rupture. Intracellular molecules may act as danger-associated molecular patterns (DAMPs) or alarmins upon release in the extracellular space. DAMPs are host-derived molecules that engage pattern recognition receptors (PRRs) in the context of cell damage or stress. This can be due to the release of molecules from certain circumscribed compartments into compartments that are normally devoid of these molecules (e.g. mitochondrial DNA into the cytoplasm) or the modification of endogenous molecules that render them PRR-agonistic (e.g. oxidized lipids). Conceptually, DAMP-dependent PRR activation is a trade-off of the sensitivity of the PRR system and not intentional. Alarmins, on the other hand, are pre-formed molecules that are meant to engage pro-inflammatory signaling cascades upon extensive cell or tissue damage. This, for example, includes certain cytokines of the IL-1 family that are sequestered in the nucleus, lacking a secretion signal (e.g. IL-1α or IL-33) ^22^. These molecules are only released upon lytic cell death, engaging their dedicated cytokine receptors.

Evidences of a contribution of necroptosis to inflammatory diseases comprise abnormal expression of RIPK3, RIPK1 and MLKL observed in samples from Crohn’s Disease patients, suggesting a role of necroptosis in epithelial cell death and inflammation of the terminal ileum ^23^. In addition, homozygous mutations in the CASP8 gene leading to caspase-8 deficiency and increased necroptosis have been described in patients with very early onset inflammatory bowel disease (VEO-IBD) ^24^. Moreover, necroptosis has been implicated in the development of neurodegenerative and cardiovascular diseases, as well as in different liver disorders ^20,25^. However, the complexity of these pathologies makes it difficult to establish necroptosis as a direct driver of inflammation. *In vivo* studies in Ripk3 or Mlkl deficient mice have been conducted to explore the role of necroptosis in different disease models. Nevertheless, discordant conclusions have been reported regarding the importance of necroptosis as an inducer of inflammation ^20,26^. Despite the number of studies on necroptosis-mediated inflammation, the identity of molecules that may act as DAMPs or alarmins upon release by necroptotic cells remains elusive. Multiple reasons account for the difficulties in establishing a causative link between putative danger signals and necroptosis-mediated inflammation, including the use of immune stimuli to induce necroptosis, the necroptosis-independent inflammatory roles of RIPK3 ^27^ and the complexity of the investigated *in vivo* models ^20^. In light of these complications, we here sought to establish an *in vitro* system to explore the identity of necroptosis-associated DAMPs/alarmins that may be implicated in necroptosis-driven inflammation.

## Results

### A genetic system to study necroptosis-dependent inflammatory signaling

To study how necroptosis results in the release of pro-inflammatory mediators *in vitro,* we turned to BLaER1 monocytes as a model system. In the absence of caspase-8, these cells display a strong necroptotic phenotype following TLR4 stimulation ^13^. To this end, we employed *CASP8*^-/-^ cells as necroptosis-prone cells and *CASP8*^-/-^ × *MLKL*^-/-^ cells as necroptosis resistant controls in a *CASP4*^-/-^ genetic background, to prevent possible non-canonical inflammasome activation upon LPS stimulation (Fig. S1A). Under these conditions, necroptosis induction occurs rapidly as evidenced by measuring LDH release in the supernatant (Fig. 1A and Fig. S1B) or when assessing the phosphorylation status of MLKL (Fig. 1B). In order to address whether necroptotic cells can trigger inflammatory responses in bystander cells, we devised a system in which we could co-culture cells undergoing necroptosis (donor cells) with cells that were insensitive to necroptosis induction (recipient cells). In this setting, we measured interleukin-6 (IL-6) production in recipient cells as a proxy for pro-inflammatory gene expression (Fig. 1C). IL-6 was chosen for its high sensitivity and dynamic range as an NF-κB target gene. The *IL6* gene was deleted in the donor cell population, so that all IL-6 measured in this setup can be ascribed to recipient cells. Apart from its intended phenotype, IL-6 deficiency had no impact on necroptosis or pro-inflammatory gene expression in donor cells (Fig. S1C-E). Further, since LPS is used in this model to trigger necroptosis in donor cells, we rendered recipient cells insensitive to LPS by disrupted the *TLR4* gene (Fig. 1C). Donor and recipient cells were then co-cultured and analyzed for LDH release, as well as IL-6 production. As expected, co-cultures containing *CASP8*^-/-^ donor cells that were additionally deficient for IL-6 displayed a marked release of LDH into the supernatant upon LPS treatment (Fig. 1D, left panel). This response was not seen when *CASP8*^-/-^ × *MLKL*^-/-^ donor cells were used. Intriguingly, when co-cultured with necroptosing donor cells but not *CASP8*^-/-^ × *MLKL*^-/-^ cells, a strong IL-6 response was seen in recipient cells (Fig. 1D, right panel). To explore whether this pro-inflammatory activity of necroptosis required cell-to-cell contact, we next conducted experiments, in which we transferred supernatant from necroptosing donor cells or control conditions onto recipient cells. Analogous to the co-culture conditions, a potent IL-6 response was seen when supernatant from stimulated *CASP8*^-/-^ cells was transferred (Fig. 1E). In line with these results, transfer of supernatant from necroptosing donor cells led to NF-κB activation in recipient cells as evidenced by p65 and IκBα phosphorylation, as well as IκBα degradation (Fig. 1F). Altogether, these results indicated that necroptosis-driven inflammatory responses can be recapitulated in an *in vitro* system, and that cell-to-cell contact is not required to evoke pro-inflammatory activity.

**Fig. 1.**
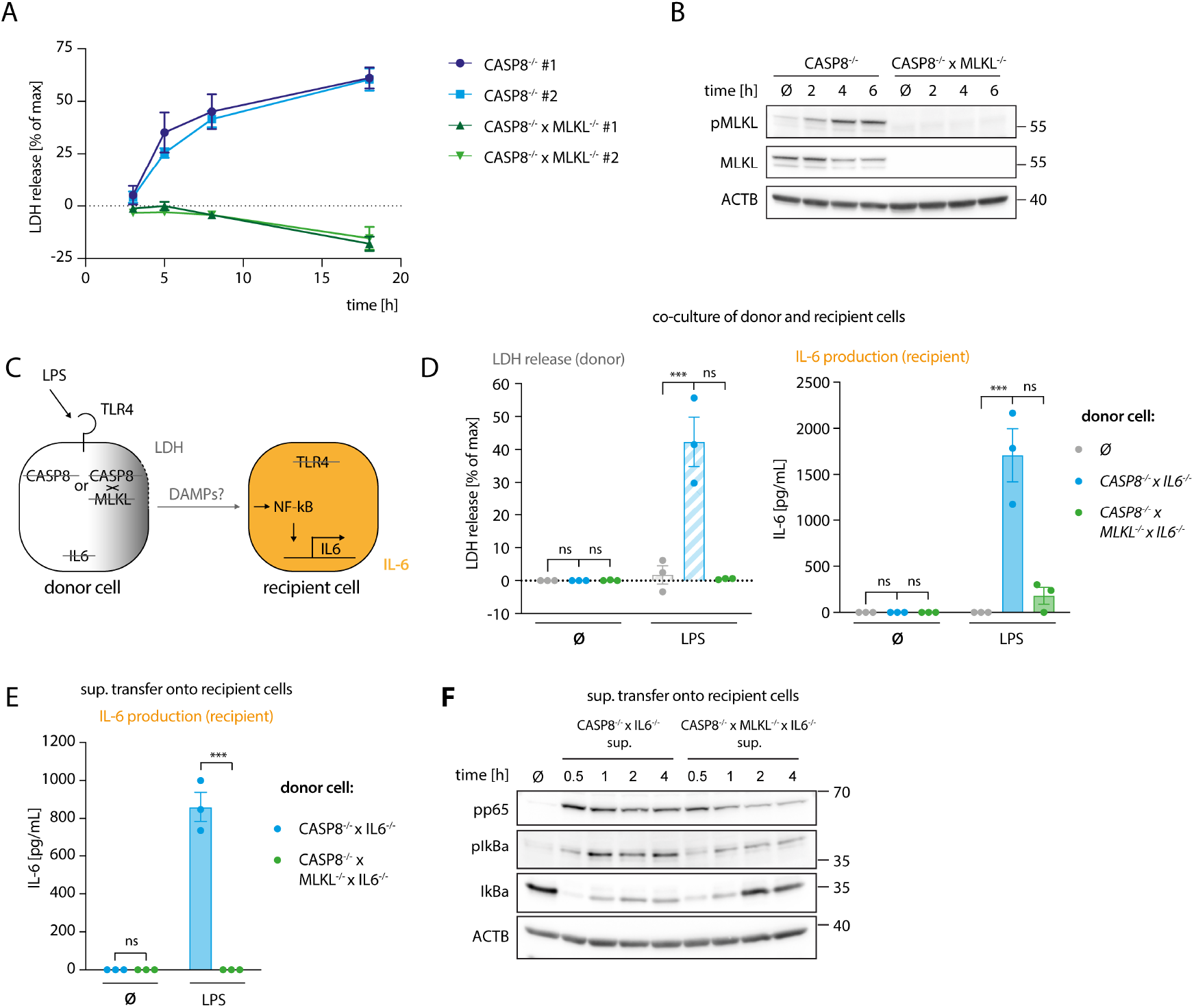
A genetic system to study necroptosis-dependent inflammation. (A) BLaER1 cells of the indicated genotypes were stimulated with 2 ng/ml LPS for 3 h, 5 h, 8 h or 18 h and LDH release was determined. (B) *CASP8*^-/-^ and *CASP8*^-/-^ × *MLKL*^-/-^ BLaER1 macrophages were stimulated with 2 ng/ml LPS for the indicated times. Cell lysates were blotted for the indicated proteins. (C) Schematic overview of the system of donor and recipient BLaER1 cells generated to perform co-culture and supernatant transfer experiments. (D) Cells of the indicated genotypes were co-cultured in a 1:1 ratio with TLR4^-/-^ recipient cells (grey dots = recipient cells only) and stimulated with 2 ng/ml LPS for 18 h. LDH release and IL-6 production were determined. (E) Donor cells were stimulated with 2 ng/ml of LPS for 18 h. The supernatant was then collected and used to stimulate TLR4^-/-^ recipient cells for 24 h and IL-6 secretion was determined. (F) *CASP8*^-/-^ × *IL6*^-/-^ and *CASP8*^-/-^ × *MLKL*^-/-^ × *IL6*^-/-^ cells were stimulated with 2 ng/ml LPS for 18 h and the supernatant was transferred on *TLR4*^-/-^ recipient cells for the indicated period of times. Immunoblotting of the indicated proteins was performed on the lysates of stimulated TLR4^-/-^ cells. Data are depicted as mean ± SEM of 3 independent experiments (A, D, and E) or as one representative experiment of two independent experiments (B, F). Statistics indicates significance by two-way ANOVA (D and E) with either a Dunnet (D) or Šidák (E) correction for multiple testing: *** P<0.001, ns=not significant. Raw data are available in Figure 1—source data 1 and 2.

### Necroptosis-driven inflammatory responses are mediated by TNF

Our results indicated that necroptosis-driven inflammation in bystander cells is largely mediated by a soluble factor. Further, considering the experimental setup that made use of *TRL4*^-/-^ recipient cells, we could exclude a TLR4-agonistic molecule as the mediator of this response. Prolonged incubation at 75°C resulted in a complete loss of the pro-inflammatory activity of the necroptotic supernatant, indicating that a proteinaceous component might play a critical role in its activity (Fig. S2A). To explore this possibility, we aimed at separating the supernatant of necroptotic cells by using size exclusion chromatography (SEC). To obtain a large volume of necroptotic supernatant, we switched to THP-1 cells as donor cells. We engineered these cells to undergo necroptosis upon expression of a dominant active MLKL construct (MLKL^1-201^) that is sufficient to induce cell death (Fig. S2B) ^28^. Such engineered cells underwent necroptosis as expected and upon transfer, the supernatant of these cells triggered IL-6 production in recipient cells (Fig. S2B and C). We then subjected supernatant of control or necroptosing THP-1 cells to SEC and tested the individual fractions for pro-inflammatory activity on recipient cells (Fig. 2A-E). These experiments revealed that the IL-6 inducing activity eluted at around 15 - 17 ml of the column volume, which corresponds to proteins of approximately 20-70 kDa in size. As a complementary approach, we also generated recipient cells, in which we deleted essential signaling hubs for various PRR systems. To block TLRs and cytokines of the IL-1 family, we deleted *MYD88* and *TICAM1* (TRIF), to prevent RLR and cGAS-dependent activation, we deleted *MAVS* and *STING1* (STING), and to block TNF and Lymphotoxin-α (LTA)-dependent pro-inflammatory gene expression we deleted *TNFRSF1A* (TNFR1) (Fig. 2F, S2D). Transferring necroptotic supernatant onto these cells in the presence of the TLR4 inhibitor CLI095 to inhibit LPS-driven effects, revealed that *TNFRSF1A*^-/-^ cells showed a markedly decreased IL-6 response, whereas the perturbation of the other signaling hubs had no impact (Fig. 2G). MyD88/TRIF and MAVS/STING deficient cells used in this assay were produced in a wildtype (WT) background, while TLR4 and TNFR1 deficient cells were generated from caspase-4-deficient cells. A such, we additionally conducted the supernatant transfer experiment to compare WT and *CASP4*^-/-^ cells in their response to the necroptotic supernatant. Caspase-4 deficiency did not account for a different behavior of these recipient cells in our supernatant transfer assay (Fig. S2E). To elucidate whether this TNFR1-dependent activity required either TNF, LTA, or both, we generated donor cells, in which these cytokines were deleted (Fig. 2H). None of these perturbations impacted on necroptosis induction (Fig. 2H, left panel). However, TNF-deficient donor cells failed to trigger IL-6 production in recipient cells, whereas LTA deficiency had no impact (Fig. 2H, middle and right panel). Since these results pointed to TNF as the main driver of pro-inflammatory gene expression in bystander cells, we next assessed TNF production of the donor cell population as a function of necroptosis induction. These experiments revealed that TNF secretion was markedly increased in cells that underwent necroptosis, with peak levels being reached already 4 hours after stimulation (Fig. 2I). Importantly, TNF release by LPS-stimulated *CASP8*^-/-^ cells could be reduced by treatment with the RIPK3 inhibitor GSK’872, ruling out that clonal differences may account for the divergence in TNF secretion between *CASP8*^-/-^ and *CASP8*^-/-^ × *MLKL*^-/-^ cells (Fig. S2F). In summary, these results indicated that necroptosis-driven induction of pro-inflammatory gene expression is largely driven by TNF in this model. Moreover, these data suggested that necroptosis strongly enhanced TNF secretion.

**Fig. 2.**
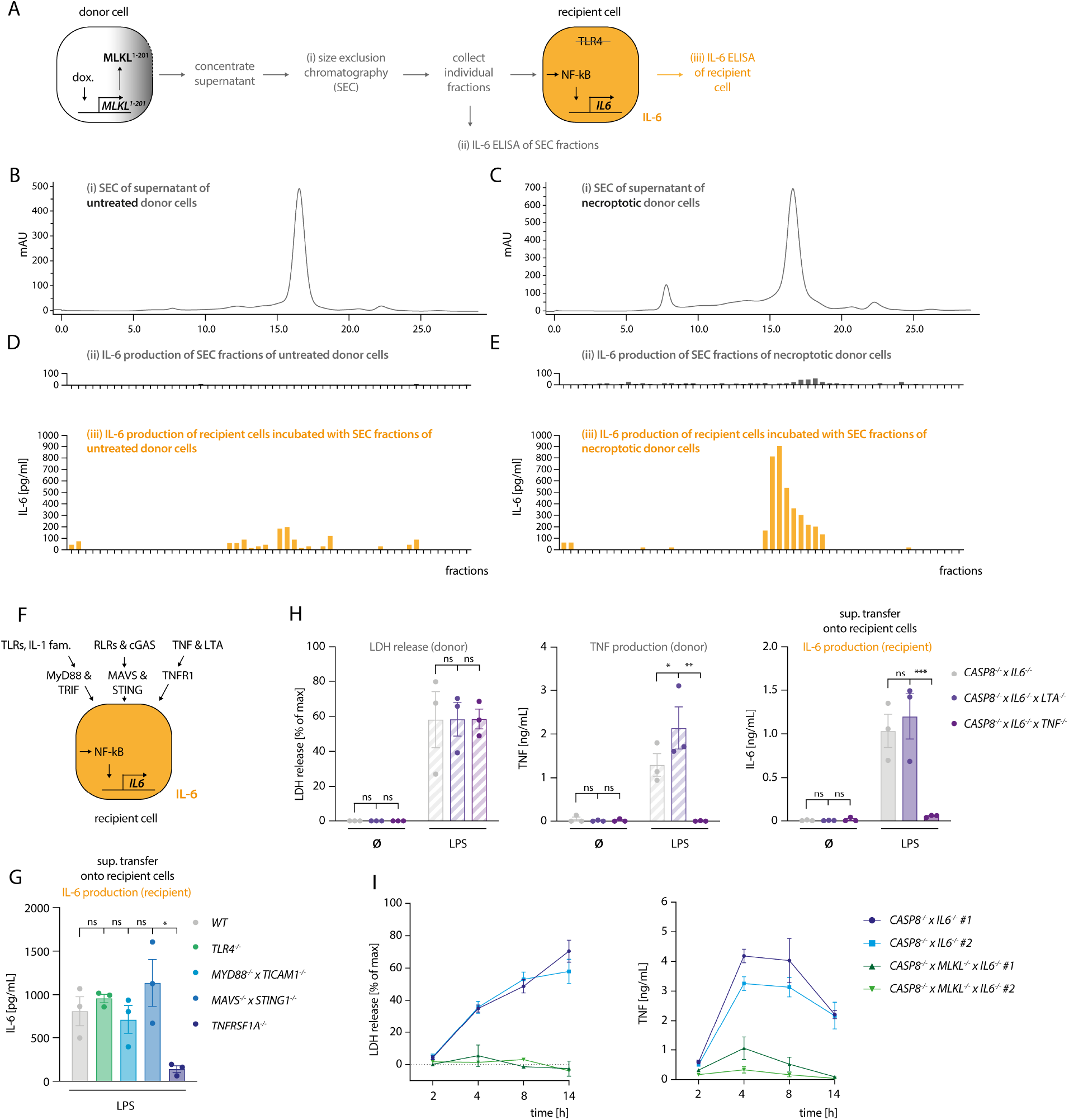
Necroptosis-driven inflammatory responses are mediated by TNF. (A) Schematic overview of the SEC experiment. (B, C) Chromatograms of SEC on the supernatant of THP1 MLKL^1-201^ cells left untreated (B) or stimulated with 1 μg/ml doxycycline for 16 h (C). (D, E) IL-6 measurement in SEC fractions of untreated and treated THP1 MLKL^1-201^ cells (in grey, upper graphs) and IL-6 production of *TLR4*^-/-^ BLaER1 recipient cells after incubation with SEC fractions for 14 h (in orange, lower graphs). (F) Scheme of knock-out generation in BLaER1 recipient cells to invalidate specified innate immune pathways. BLaER1 cells deficient for the indicated pathways were subsequently tested in the supernatant transfer assay. (G) Recipient cells of the indicated genotypes were incubated for 24 h with the supernatant of *CASP8*^-/-^ × *IL6*^-/-^ donor cells stimulated for 18 h with 2 ng/ml LPS. To block LPS-dependent effects, the TLR4 inhibitor CLI095 was added to recipient cells at the concentration of 1 μg/ml. IL-6 production of recipient cells was measured. (H) BLaER1 cells of the indicated genotypes were stimulated with 2 ng/ml LPS or left untreated for 18 h, after which LDH and TNF release were measured. The supernatant of these cells was utilized to stimulate *TLR4*^-/-^ BLaER1 cells. IL-6 release by *TLR4*^-/-^ cells was measured 24 h after. (I) BLaER1 cells of the indicated genotypes were stimulated with 2 ng/ml LPS for the indicated time points. LDH release and TNF secretion from donor cells of the specified genotypes were measured at the indicated time points. Data are depicted as mean ± SEM of 3 independent experiments (G, H and I) or as one representative experiment of two independent experiments (B-E). Statistics indicates significance by one-way (G) or two-way ANOVA (H) with a Dunnet (G and H) correction for multiple testing: *** P<0.001, ** P<0.01, * P<0.05, ns=not significant. Raw data are available in Figure 2—source data 1.

### Necroptosis boosts TNF translation and enhances TNF shedding

As a key pro-inflammatory mediator, TNF is controlled at multiple levels. Next to being regulated at the level of transcription, the translation of TNF mRNA is heavily controlled at the post-transcriptional level. To this end, several cis-elements within the 3’ UTR of TNF have been identified that are regulated by various trans-activating factors that repress TNF mRNA translation or induce decay of its mRNA ^29–33^. Moreover, the availability of the protein itself in the extracellular *milieu* is also regulated. As such, being produced as a type II transmembrane protein, TNF needs to be cleaved by membrane metalloproteases to be released. This step is mainly regulated by ADAM17 and to a lesser extent by ADAM10 ^34–37^. Comparing *CASP8*^-/-^ cells with *CASP*^-/-^ × *MLKL*^-/-^ revealed no significant difference in TNF mRNA expression between these two cell types (Fig. 3A), thereby excluding the possibility of enhanced TNF mRNA transcription. However, when we measured TNF protein expression, we observed both higher amounts of pro-TNF levels in cell lysates as well as increased levels of cleaved TNF in the supernatant, suggesting that regulation of TNF expression and shedding may both contribute to enhanced TNF release (Fig. 3B). Of note, *CASP8*^-/-^ and *CASP8*^-/-^ × *MLKL*^-/-^ cells produced similar amounts of IL-6 and IL-8, showing that other pro-inflammatory cytokines transactivated by NF-κB were not affected by these genotypes. As such, both IL-6, as well as IL-8 levels did not show any differences between *CASP8*^-/-^ and *CASP8*^-/-^ × *MLKL*^-/-^ cells (Fig. S1C, D). Speculating that necroptosis impacted on the post-transcriptional regulation of TNF mRNA translation via its 3’ UTR, we generated *CASP8*^-/-^ and *CASP8*^-/-^ × *MLKL*^-/-^ cells, in which we deleted the 3’UTR of the TNF gene (TNF Δ 3’UTR) (Fig. 3C). Doing so, we eliminated all critical cis-controlling elements that negatively affect the stability or translation of TNF mRNA, while we left the poly adenylation signal intact. As a result, pro-TNF expression levels were dramatically increased in both *CASP8*^-/-^ and *CASP8*^-/-^ × *MLKL*^-/-^ cells and considerable amounts of pro-TNF expression were already observed in lysates of unstimulated cells (Fig. 3D). Importantly, removing the 3’ UTR resulted in similar expression levels of pro-TNF for both necroptotic and non-necroptotic cells (Fig. 3D), indicating that necroptosis indeed impacts on the negative regulation of TNF mRNA translation through its 3’ UTR. In this setting, we could now investigate the impact of necroptosis on TNF shedding. Indeed, despite similar expression of pro-TNF, *CASP8*^-/-^ cells still exhibited 3-4× higher levels of soluble TNF in the supernatant upon necroptosis induction (Fig. 3D-F). Altogether, these results suggested that increased secretion of mature TNF during necroptosis depends on higher pro-TNF expression, by alleviating negative regulatory pathways at the posttranscriptional level, as well as enhanced TNF shedding.

**Fig. 3.**
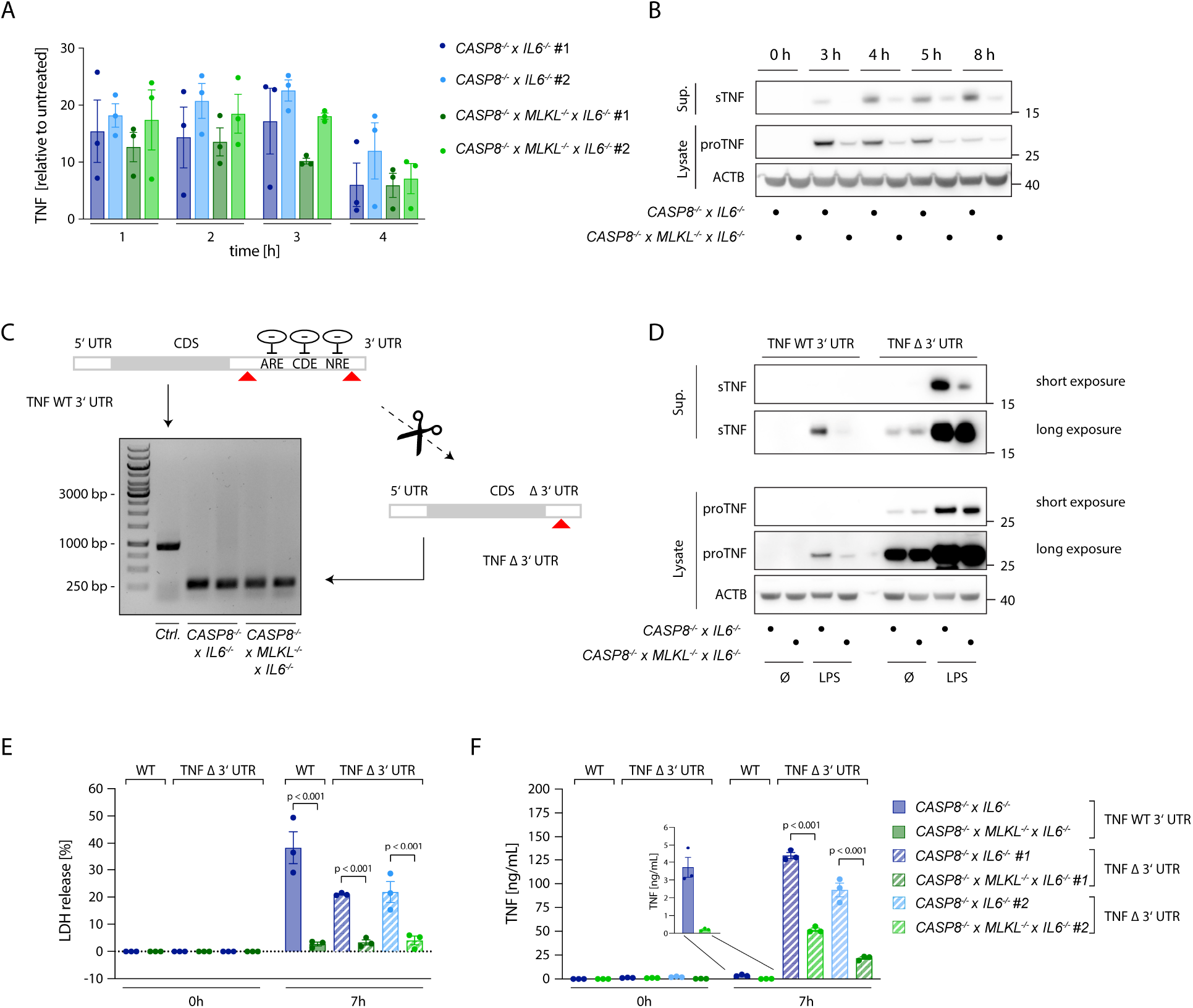
Necroptosis boosts TNF translation. (A) qPCR of TNF induced upon stimulation with 2 ng/ml LPS in the indicated cells. Results are shown as fold change relative to unstimulated cells. (B) Immunoblotting of pro-TNF (cell lysates) and sTNF (supernatants) of *CASP8*^-/-^ × *IL6*^-/-^ and *CASP8*^-/-^ × *MLKL*^-/-^ × *IL6*^-/-^ cells upon stimulation with 2 ng/ml LPS for the indicated time points. ACTB was detected as loading control. (C) *CASP8*^-/-^ and *CASP8*^-/-^ × *MLKL*^-/-^ BLaER1 cells deficient for the 3’UTR of the TNF gene (TNF Δ3’UTR) were generated by employing two gRNAs targeting the 3’UTR at its very beginning and very end, respectively (red arrows). The removal of the 3’UTR was assessed by PCR (Control = long PCR product, TNF Δ3’UTR = short PCR product). Two clones of TNF Δ3’UTR are shown per parental genotype. (D) BLaER1 cells of the indicated genotypes were stimulated with LPS (2 ng/mL) for 7 h. pro-TNF expression and matured TNF were assessed by western blotting. (E, F) BLaER1 cells of the indicated genotypes were stimulated with LPS (2 ng/mL) for 7 h. Cell death was evaluated by LDH assay (E). Secreted TNF was measured by ELISA (F). Data are depicted as mean ± SEM of three independent experiments or as one representative immunoblot of three independent experiments (B, D). Statistics indicates significance by two-way ANOVA with a Tukey correction for multiple testing: *** P<0.001. Raw data are available in Figure 3—source data 1, 2 and 3.

### Necroptosis enhances TNF shedding

To explore how necroptosis impacts on TNF shedding, we used 293T cells as a heterologous expression system, in which we could dissociate TNF expression from shedding. To induce necroptosis, we expressed the autoactive MLKL^1-201^ construct, while using an inactive MLKL variant (MLKL^1-154^) or mCherry as a control ^28^. Of note, this reductionist setting could also rule out RIPK3-dependent effects on necroptosis-dependent TNF secretion. MLKL^1-201^-dependent necroptosis induction led to a marked release of LDH from 293T cells, while the other conditions showed little activity (Fig. 4A, left panel). Concomitant with cell death, a strong TNF signal in the supernatant was detected by ELISA (Fig. 4A, right panel). This increase in TNF within the supernatant was due to a markedly enhanced pro-TNF conversion to soluble TNF, as evidenced by immunoblot. Importantly, the release of IL-6, which does not require shedding for its maturation, was not enhanced upon necroptosis induction in these settings (Fig. S3A). To confirm that necroptosis indeed resulted in increased TNF shedding and not in a passive release of pro-TNF, we additionally conducted experiments in 293T cells lacking ADAM10 and ADAM17 ^38^. In these cells, TNF release upon necroptosis was severely impaired, while ADAM10/17 deficiency had no impact on cell death following MLKL^1-201^ expression (Fig. 4B). Analogous results were obtained when studying TNF levels in lysates or supernatant by immunoblot (Fig. 4B, lower right panel). Next, we sought to investigate the role of ADAM proteases in the BLaER1 cells model. RNA-seq analysis revealed that ADAM10, ADAM17 and ADAM9 are the main ADAM proteases expressed in these cells and that LPS positively modulates their expression (Fig. S3B). ADAM17 and ADAM10 being the main TNF sheddases described, we generated BLaER1 cells lacking either of these two enzymes (Fig. S3C) to address their role in necroptosis-dependent TNF shedding. Doing so revealed that ADAM17 was the predominant enzyme required for TNF shedding in the context of necroptosis, while neither ADAM17 nor ADAM10 had a significant impact on cell death induction (Fig. 4C, left and middle panel). Consistent with this notion, supernatant from *ADAM17*^-/-^ but not control or *ADAM10*^-/-^ cells displayed a markedly reduced activity in the bystander cell activation assay (Fig. 3C, right panel). Additionally, the secretion of another known target of ADAM17, IL6-Rα^39^, increased in necroptotic BLaER1 cells compared to control cells (Fig. S3D). Altogether, these results indicated that necroptosis drives enhanced TNF-dependent inflammatory responses in two ways: on the one hand, necroptosis results in enhanced TNF translation, and on the other hand, necroptosis triggers TNF shedding in a predominantly ADAM17-dependent manner.

**Fig. 4.**
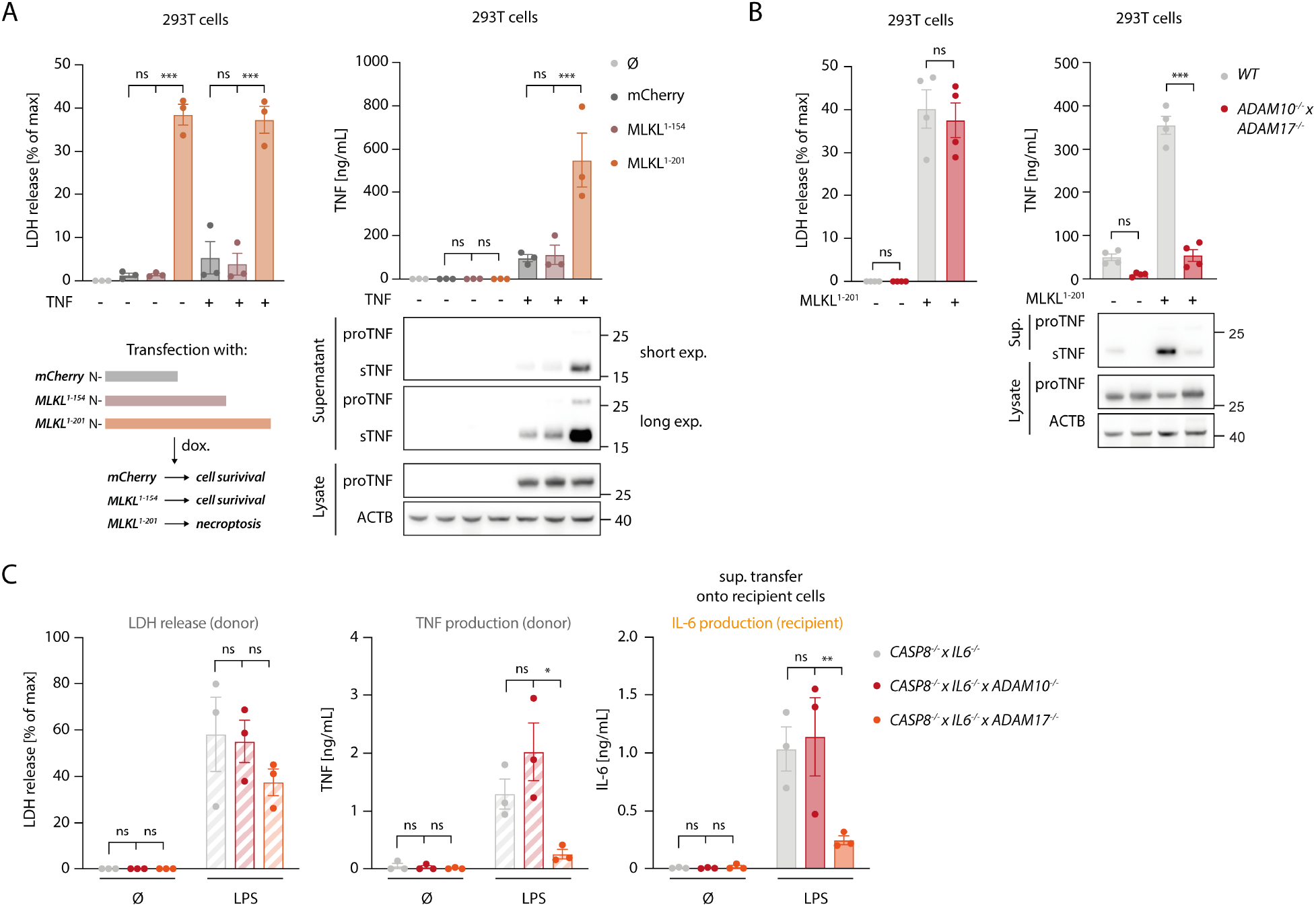
Necroptosis drives ADAM-dependent TNF shedding. (A) LDH release, TNF secretion and immunoblotting of the indicated proteins of 293T cells transfected with the specified MLKL constructs in combination with pRP-TNF (or pRP-mCherry) and stimulated the day after with doxycycline for 7 h. (B) 293T cells of the indicated genotypes were transfected with pRP-TNF and pLI-MLKL^1-201^ (or pLI-mCherry). LDH and TNF secretion were then determined. (C) BLaER1 cells of the indicated genotypes were stimulated with 2 ng/ml LPS or left untreated for 18 h. The supernatant of these cells was utilized to stimulate *TLR4*^-/-^ BLaER1 cells. For the control supernatant of *CASP8*^-/-^ × *IL6*^-/-^ cells, the same sets of data are shown as in Fig. 2H. LDH and TNF secretion from donor cells and IL-6 release by *TLR4*^-/-^ cells were measured. Data are depicted as mean ± SEM of three (A and C) or four (B) independent experiments or as one representative immunoblot of three (A, B) independent experiments. Statistics indicates significance by two-way ANOVA (A) with a Dunnet (A and C) or Šidák (B) correction for multiple testing: *** P<0.001, ** P<0.01, * P<0.05, ns=not significant. Raw data are available in Figure 4—source data 1 and 2.

### Necroptosis triggers shedding activity in a switch-like fashion

To explore the spatiotemporal relationship of necroptosis and TNF shedding, we made use of a TNF shedding reporter construct (C-tag TNF) that allows the monitoring of TNF maturation at the ADAM17 cleavage site in real time ^38^. Upon cleavage, this construct exposes a neo C-terminus that serves as an epitope for a nanobody that only binds the cleaved version of this reporter. This construct was additionally equipped with an N-terminal mCherry tag, to monitor the subcellular localization of TNF. Cells expressing the C-tag reporter displayed the same gain in TNF shedding upon necroptosis induction as cells expressing unmodified TNF (Fig. S4A). Using a fluorescently labeled nanobody, we monitored the C-tag signal as a function of MLKL^1-201^-induced necroptosis in control cells, as well as in *ADAM10*^-/-^ × *ADAM17*^-/-^ deficient cells. In addition, we also measured C-tag positivity in controls not undergoing necroptosis. 4.5 hours following necroptosis induction, we observed a strong signal for the C-tag nanobody at the plasma membrane of cells that also showed signs of necroptosis (Fig. 5A and 5B, green channel). This signal was completely abrogated in cells lacking ADAM10/17, which still displayed membrane-associated expression of TNF (Fig. 5A, red channel). Control cells not undergoing necroptosis, also showed a C-tag reactive signal, yet with marked lower intensity. Time-lapse analysis revealed that cells undergoing necroptosis became positive for the C-tag signal within 30-40 minutes (Fig. 5B and C). Quantifying the C-tag signal and analyzing the slope for several cells individually revealed that cells undergoing necroptosis indeed displayed a rapid, switch-like gain in TNF shedding. On the other hand, the gain in C-tag signal for cells not undergoing necroptosis was only gradual, and it never reached the same level as necroptosing cells in the time frame studied (Fig. 5B and D). Similar to apoptosis, necroptosis also results in the externalization of phosphatidylserine phospholipids to the outer leaflet of the membrane ^40^. Measuring Annexin V positivity next to TNF shedding, we observed that these two events were closely correlated, with the individual slopes of the gain in fluorescent signal being comparable (Fig. S4B-D). However, C-tag positivity preceded Annexin V positivity by a mean value of 24 minutes (Fig. S4E). Studying PI positivity as a proxy for membrane permeabilization, we observed that cells undergoing necroptosis and TNF shedding displayed a delayed gain in PI signal (Fig. S4F-I). In comparison to Annexin V, this delay was even more pronounced with a mean difference of 53 minutes (Fig. S4I). Further, analyzing the slope of the gain in PI positivity revealed that this event occurred more rapidly than the TNF shedding event (Fig. S4G and H). This was also reflected in the slope of the PI signal which was approximately twice the value of the C-tag signal (Fig. S4H). Altogether, these results indicated that necroptosis-induced TNF shedding is a rapid event that differs from the steady state shedding process in magnitude and kinetics. As such, it displays a switch-like behavior, in which TNF shedding occurs in a time frame of 30-40 minutes. In the course of cell death, TNF shedding precedes the outward flipping of phosphatidylserine phospholipids, which is followed by membrane permeabilization.

**Fig. 5.**
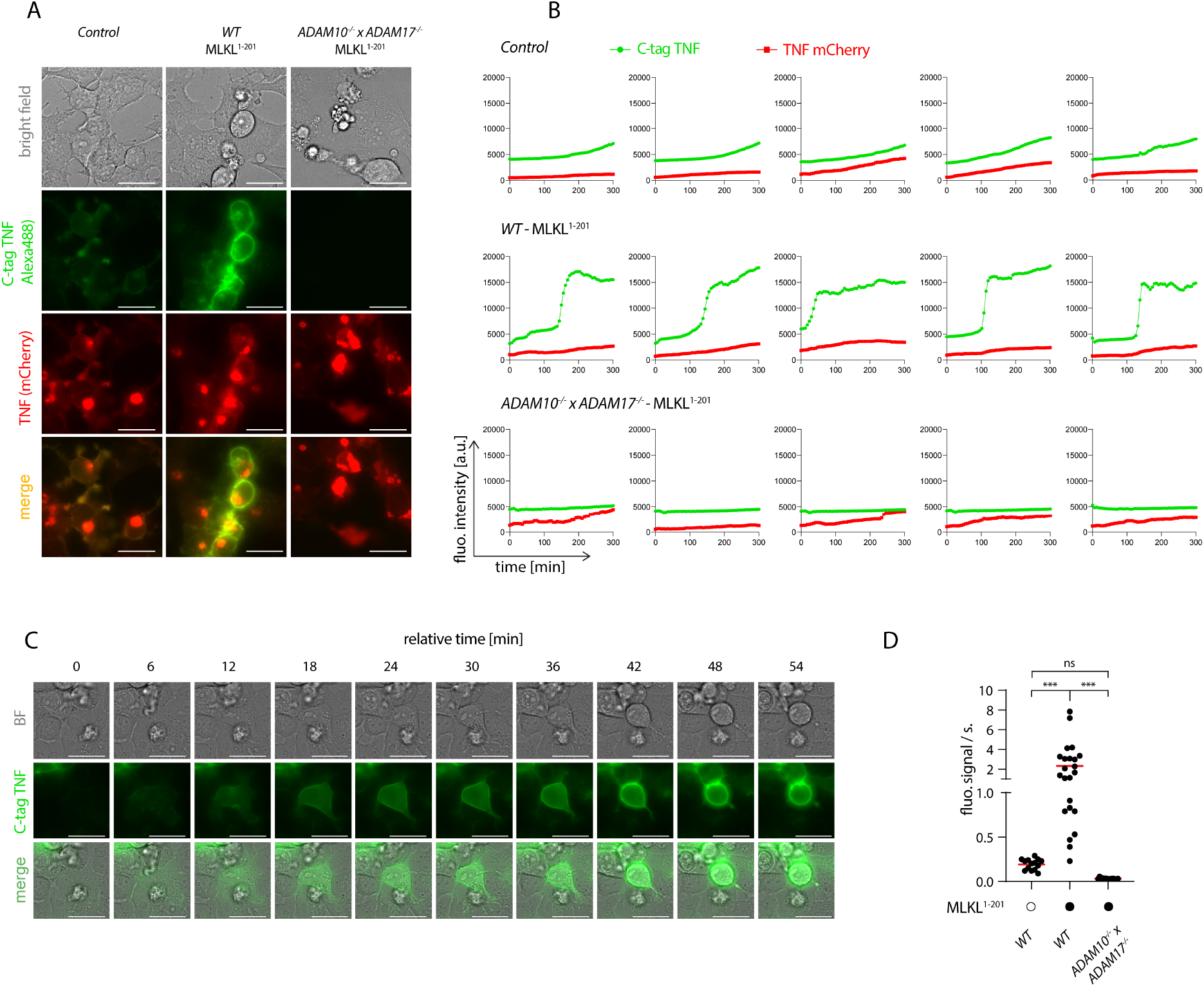
Necroptosis triggers rapid shedding activity. (A) 293T (control), 293T MLKL^1-201^ and *ADAM10*^-/-^ × *ADAM17*^-/-^ MLKL^1-201^ 293T were transfected with pLI-C-tag-TNF linker mCherry and pEF-BOS-nBFP and stimulated with 1 μg/ml doxycycline and imaged over time starting 4 h after doxycycline treatment. Exemplary images of the indicated channels 8h30 after doxycycline induction are shown. Bar = 25 μm. (B) Trajectories of TNF mCherry and C-tag TNF of cells treated as in (A). Five representative cells per cell type are shown. (C) One exemplary cell from the 293T MLKL^1-201^ population stimulated as in (A) is shown. Bar = 25 μm. (D) Calculated slopes of the C-tag TNF channel signal of cells stimulated as in (A) are depicted, while the mean value is highlighted in red. Representative data of three (A and C) independent experiments are shown. Statistics indicates significance by one-way ANOVA (D) with a Tukey correction for multiple testing: *** P<0.001, ns=not significant. Raw data are available in Figure 5—source data 1.

### Enhanced TNF shedding is a general phenomenon of lytic cell death rather than a specific necroptosis-related feature

Our data thus far indicated that necroptosis triggers both TNF translation and shedding and that the latter takes place in the early stages of cell death, before massive membrane damage. Live cell imaging experiments revealed that TNF shedding occurs as a switch-like phenomenon and appears to be strictly related to physical changes of the cell membrane before membrane permeabilization. We therefore asked whether the mechanism behind increased TNF shedding is specifically related to the necroptotic pathway or if it is promoted by changes in membrane integrity during lytic cell death in general. To address this question, we applied a model in which we can induce lytic cell death by adding an exogenous pore-forming molecule-namely, the pore-forming toxin streptolysin O (SLO) *from S. pyogenes* ^41,42^. As for the MLKL gain-of-function studies, we used HEK 293T cells overexpressing pro-TNF that we then challenged with exogenous SLO. As expected, SLO treatment led to lytic cell death, as indicated by LDH release (Fig. 6A, left panel). Similar to the overexpression of MLKL^1-201^, SLO-mediated cell death was accompanied by massive TNF secretion due to enhanced shedding, as evidenced by ELISA and immunoblot (Fig. 6A, right panel and Fig. 6B). Importantly, IL-6 secretion was not enhanced upon SLO-mediated cell death, but rather decreased due to cell death during the experiment (Fig. 6C). In conclusion, these data indicate that enhanced TNF shedding is linked to the membrane dynamics upon lytic cell death rather than being a specific feature of the necroptotic pathway itself.

**Fig. 6.**
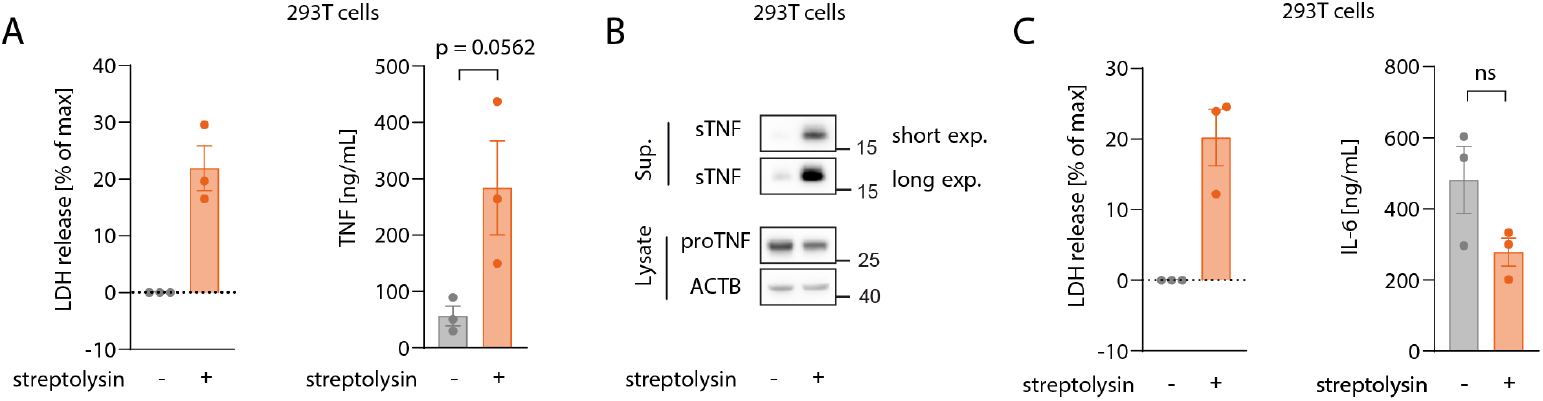
Enhanced TNF shedding is a general phenomenon of lytic cell death rather than a specific necroptosis-related feature. (A and B) 293T cells were transfected with 100 ng/well of pRP_TNF in a 96 well format. After 18h, medium was changed with or without streptolysin for 7 h. Supernatant and cell lysates were collected for LDH, ELISA (A) and immunoblot (B). (C) 293T cells were transfected with 100 ng/well of pRP_IL6 in a 96 well format and treated as in (A). LDH release and IL-6 secretion were assessed in the supernatant. Data are depicted as mean ± SEM of three independent experiments. Statistics (A and C) indicates significance by an unpaired two-tailed t-test: P value as indicated or ns=not significant. Raw data are available in Figure 6—source data 1 and 2.

### Necroptosis-driven TNF release in primary cells

Thus far our experiments had relied on cell lines to study the relevance and mechanism of necroptosis-induced TNF shedding. To explore whether these events also play a role in primary cells undergoing necroptosis, we turned to primary, human monocyte-derived macrophages (hMDM) as a model. Treating these cells with LPS and the pan-caspase inhibitor zVAD, resulted in a lytic cell death in approximately 25% of the cell population within an 8-hour timeframe (Fig. 7A, left panel). Treatment with the specific RIPK3 inhibitor GSK’872 blocked this cell lysis, indicating that cells died by necroptosis. Analyzing TNF secretion revealed that necroptosis induction doubled the amount of released TNF, while it decreased IL-6 production by half (Fig. 7A, middle and right panel). To employ a system that does not require LPS stimulation to induce necroptosis, we used murine macrophages derived from hematopoietic stem cells (HSCs). As such, we transduced HSCs with a murine RIPK3 construct that can be dimerized to induce necroptosis upon the addition of the small molecule drug AP20187 (Fig. 7B) ^43^. After differentiation of these HSCs, these cells displayed a typical macrophage-like morphology and about 30% of them expressed the RIPK3 dimerizing construct, as judged by EGFP expression (Fig. S5A and S5B). Indeed, RIPK3 dimerization by treatment with AP20187 induced lytic cell death specifically in transduced cells. Moreover, this synthetic necroptosis-induction resulted in TNF release of these macrophages in the supernatant (Fig. 7C). In summary, these results suggested that primary cells undergoing necroptosis also display enhanced TNF secretion.

**Fig. 7.**
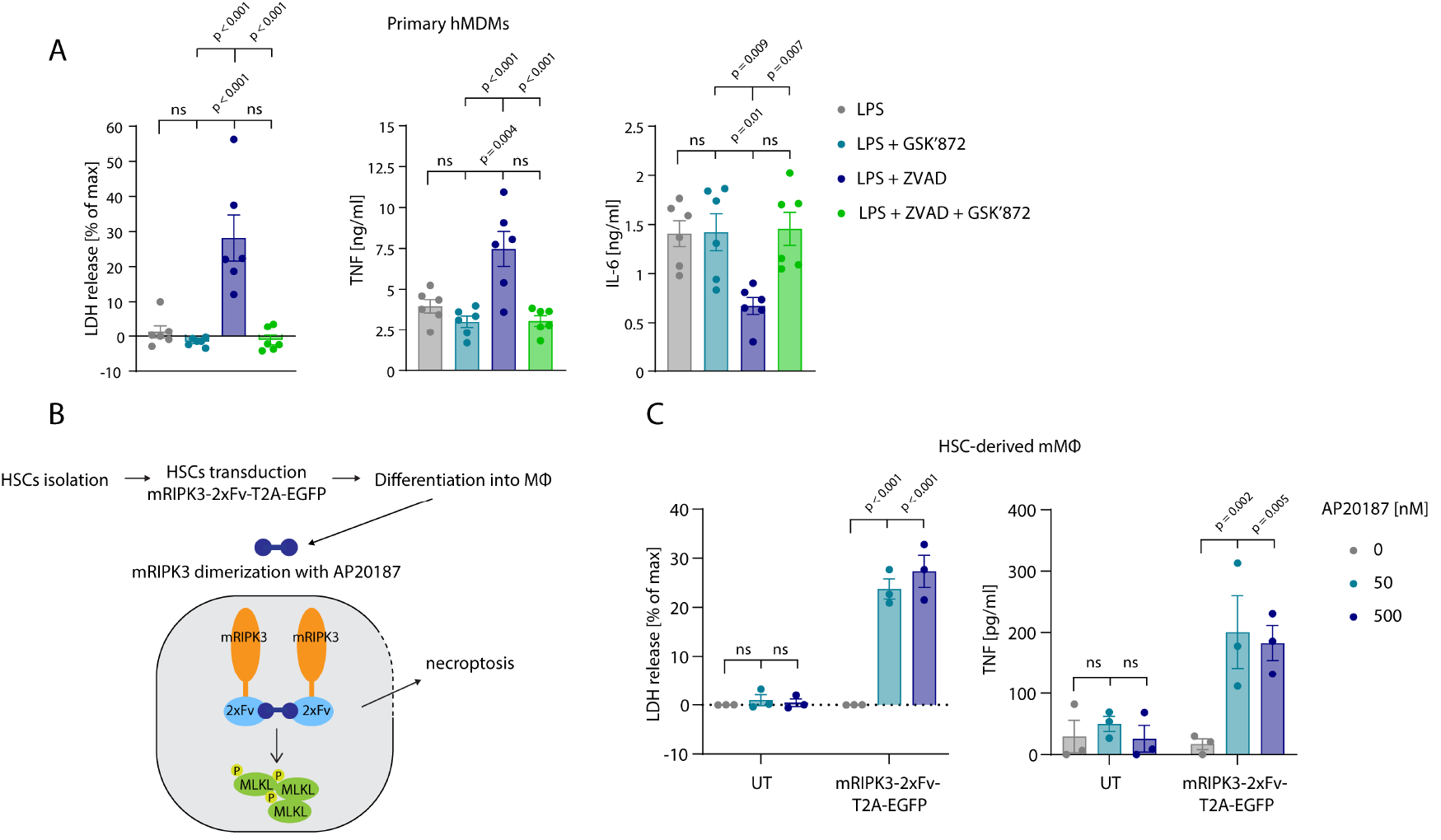
Necroptosis driven TNF release in primary cells. (A) Primary hMDMs were stimulated for 8 h with 0.2 ng/ml LPS and the specified inhibitors (GSK’872 = 3μM, ZVAD = 20 μM). LDH release and cytokine production were measured (B) Schematic overview of experimental set-up with HSC-derived macrophages. (C) HSC-derived macrophages left untransduced (UT) or transduced with pFUGW-mRIPK3-2xFv-T2A-EGFP were stimulated with the indicated amount of AP20187 for 15 h. LDH and TNF release were measured. Data are depicted as mean ± SEM of six different donors (A) or of three independent experiments (C). Statistics indicates significance by one-way ANOVA (A) or two-way ANOVA (C) with a Tukey (A) or Dunnet (C) correction for multiple testing: *** P<0.001, ** P<0.01, * P<0.05, ns=not significant. Raw data are available in Figure 7—source data 1.

## Discussion

Necroptosis has emerged as a highly inflammatory form of regulated necrotic cell death. Numerous studies, primarily in genetic mouse models, provided experimental evidence that necroptotic cell death induces inflammation *in vivo*, suggesting that necroptosis may also be involved in the pathogenesis of human inflammatory diseases. However, the mechanisms by which necroptotic cells trigger inflammatory responses in the tissue microenvironment remain poorly understood. While genetic *in vivo* models are well-suited to characterize the impact of individual candidate factors on the organismic and tissue level, it is laborious and technically difficult to establish mechanistic relationships between putative DAMP or alarmin molecules being released and their pro-inflammatory activity. A major difficulty for such ventures is the dissection of the cellular source and target for putative inflammatory signals in a biologically redundant and dynamic system. Moreover, it is inherently challenging to discover and characterize novel DAMPs and/or alarmin(s) in the *in vivo* context.

We therefore developed a reductionist *in vitro* setting, in which necroptosis-driven inflammatory pathways can be addressed by genetically manipulating donor and recipient cells. To do so, we employed a myeloid cell system, in which we could recapitulate the phenomenon of necroptosis-driven inflammation. In this setup, we observed a strong, necroptosis-dependent NF-κB response in recipient cells when either co-cultured with necroptosing cells or exposed to their supernatant. The pro-inflammatory activity of this supernatant could be further narrowed down to be a heat-sensitive factor that could be resolved by size exclusion chromatography. Systematic perturbation of sensing and signal transduction cascades in recipient cells provided the surprising notion that this pro-inflammatory activity could be largely ascribed to TNF. Indeed, deleting *TNF* in necroptotic cells or *TNFR1* in recipient cells, led to a complete loss of IL-6 release, indicating that NF-κB activation in this system is fully dependent on the TNF-TNFR1 axis.

TLR4-activated *CASP8*^-/-^ cells released far more TNF into the supernatant compared to their *CASP8*^-/-^ × *MLKL*^-/-^ counterparts, which could be ascribed to two mechanisms: on the one hand, cells undergoing necroptosis produced more pro-TNF at the protein level than necroptosis-resistant cells. Of note, the expression of other pro-inflammatory mediators was not increased under these conditions. Moreover, heightened pro-TNF levels were not due to increased pro-TNF expression at the mRNA level, indicating that post-transcriptional mechanisms are operational. These results therefore implied that necroptosis blocks an inhibitory mechanism that regulates TNF expression at the post-transcriptional level through its 3’ UTR. In line with this notion, LPS-stimulated cells undergoing necroptosis, yet lacking the TNF 3’UTR, showed similar amounts of pro-TNF expression as non-necroptosing cells, while TNF levels were strongly increased under these conditions. Mechanistically, it is conceivable that certain TNF 3’ UTR trans-activating factors are selectively depleted in the course of necroptosis or that certain post-translational modifications block their activity. Previous work has established that cell undergoing necroptosis indeed do not shut down protein synthesis ^44^ and that pro-inflammatory cytokine expression even proceeds in cells irreversibly committed to death upon the loss of cell membrane integrity ^45^. However, since pro-TNF requires further post-translational maturation at the plasma membrane, heightened pro-TNF expression levels by themselves would not be able to explain how necroptosis boosts the release of mature TNF. Indeed, we also observed that necroptosis strongly enhanced TNF shedding. We focused on this latter phenomenon in the following, since this mechanism likely constitutes the rate-limiting step in mature TNF secretion under these conditions. Moreover, this mechanism is universally applicable to cells undergoing necroptosis in the absence of a strong TNF-triggering stimulus such as the one used here. Indeed, cells heterologously expressing TNF displayed a strong increase in TNF maturation upon necroptosis induction, which was dependent on ADAM10/17. This is in line with previous studies that found that ADAM proteases are activated upon necroptosis ^46,47^. By studying TNF shedding in live cell imaging experiments, we observed that cells undergoing necroptosis displayed a far higher signal for TNF shedding at the single-cell level and that necroptosis-associated TNF maturation occurred with faster kinetics compared to control cells. In addition, we found that living cells shed TNF before MLKL-dependent membrane rupture. In particular, TNF shedding preceded PI uptake by almost one hour and it also occurred prior to Annexin V positivity. While these results would preclude phosphatidylserine (PS) exposure as the decisive factor for ADAM17 activation as previously reported ^48^, it cannot be excluded that PS exposure below the detection limit of our assay is of relevance.

To determine whether ADAM-dependent TNF release is a specific feature of necroptosis or applies generally to lytic types of cell death, we induced a necroptosis-independent type of lytic cell death using a bacterial pore-forming toxin. As such, we extracellularly added SLO, which assembles into large heterogeneous pores of 30–50 nm in diameter ^49^. Surpassing a critical concentration that cannot be repaired, SLO pores have been shown to trigger plasma membrane rupture and thus complete cell lysis. Indeed, employing SLO to induce plasma membrane permeability, we observed massive TNF shedding in cells heterologously expressing TNF. Therefore, we can exclude that a necroptosis-specific signaling molecule or unique aspect of its lytic cell death is required to trigger ADAM activity. We rather speculate that changes of the biophysical properties of the plasma membrane constitute the common denominator of ADAM activation, yet the exact mechanism of this phenomenon requires future studies. In this context, the involvement of protein kinase C (PKC) isoforms is likely, as this enzyme family is involved in the activation of ADAM17 ^50^.

Importantly, our findings could be reproduced in primary cell systems, also when using different necroptosis-inducing strategies. Both human monocyte-derived macrophages, as well as murine HSC-derived macrophages displayed an increase in TNF release upon necroptosis induction. Given the constitutive expression of TNF in the latter system, we could also make use of a genetic model that did not require a pro-inflammatory signal to induce TNF expression. Addressing the role of TNF in necroptosis-driven inflammation *in vivo* is difficult, since TNF also acts upstream of necroptosis induction. Indeed, several genetic models of necroptosis have shown a beneficial role for TNFR1 or TNF ablation in ameliorating necroptosis associated inflammation ^51–56^, and thus far this has been attributed to the TNF-TNFR1 axis to induce necroptosis. However, it is reasonable to speculate that ablation of TNFR1 also affects the inflammatory response in these models by preventing activation of bystander cells through necroptosis-associated release of TNF. Such a scenario would position TNF in a vicious cycle, in which TNF-triggered necroptosis can result in the release of TNF itself.

Although TNF plays an almost non-redundant role as a pro-inflammatory mediator in our system, additional DAMPs or alarmins downstream of necroptosis cannot be excluded. Indeed, being limited to the reductionist *in vitro* culture conditions tested here, our system cannot account for the complexity of an *in vivo* setting, in which multiple different cell types interact in a dynamic crosstalk. Moreover, by using IL-6 as a proxy for the inflammatory response, it should also be noted that our model was only designed to identify NF-κB-activating factors. Finally, by using LPS as a necroptosis-inducing signal, we had to delete TLR4 in recipient cells, which excludes numerous DAMPs that have been described to engage this PRR.

In summary, we established necroptotic cells as a key source of TNF secretion. Conceptually, in light of the here-uncovered mode of action, TNF qualifies as an alarmin molecule that is released upon cell damage. The success of anti-TNF therapies in diseases characterized by chronic inflammation may be partially explained by uncontrolled necroptosis enhancing TNF production. Importantly, the identification of TNF as a necroptosis-associated alarmin may provide the rationale for the future application of anti-TNF agents to treat pathologies with an established role of necroptosis as a driver of inflammation.

## Supplementary figures

**Fig. S1.**
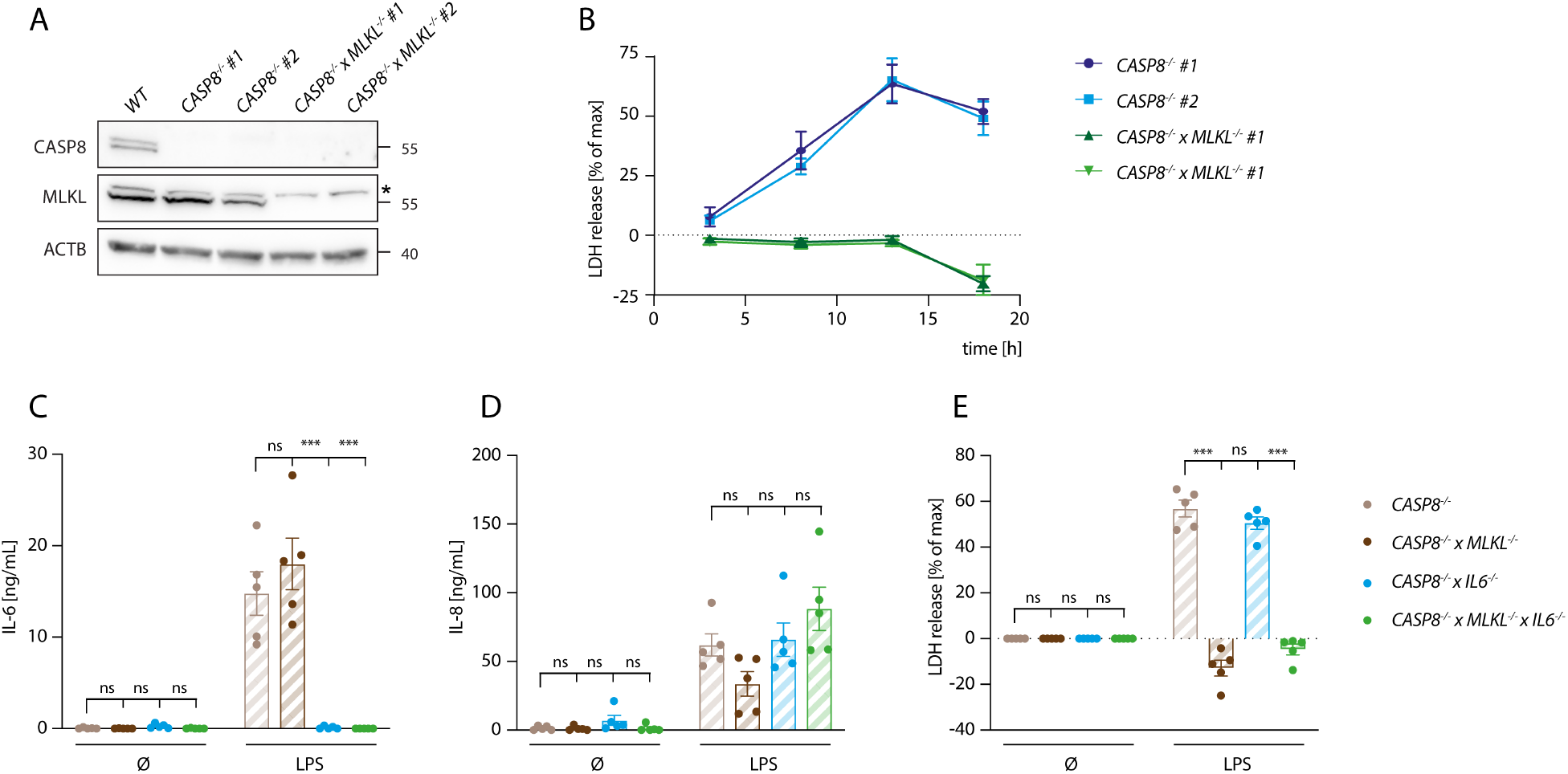
A genetic system to study necroptosis-dependent inflammation. (A) Caspase-8 and MLKL deficiency of the clones used in the study was confirmed by immunoblotting. Two clones per genotypes are shown. ACTB was detected as loading control. * denotes an unspecific band. (B) *CASP8*^-/-^ and *CASP8*^-/-^ × *MLKL*^-/-^ BLaER1 macrophages were stimulated with 200 ng/ml LPS for the indicated time points and LDH release was measured. (C, D) BLaER1 cells of indicated genotypes were stimulated with 2 ng/ml LPS or left unstimulated. IL-6 (C), IL-8 (D) and LDH release (E) were measured. Data are depicted as mean ± SEM of three (B) or five (C-E) independent experiments. Statistics indicates significance by two-way ANOVA (C-E) with a Dunnet correction for multiple testing: *** P<0.001, ns=not significant. Raw data are available in Figure S1—source data 1 and 2.

**Fig. S2.**
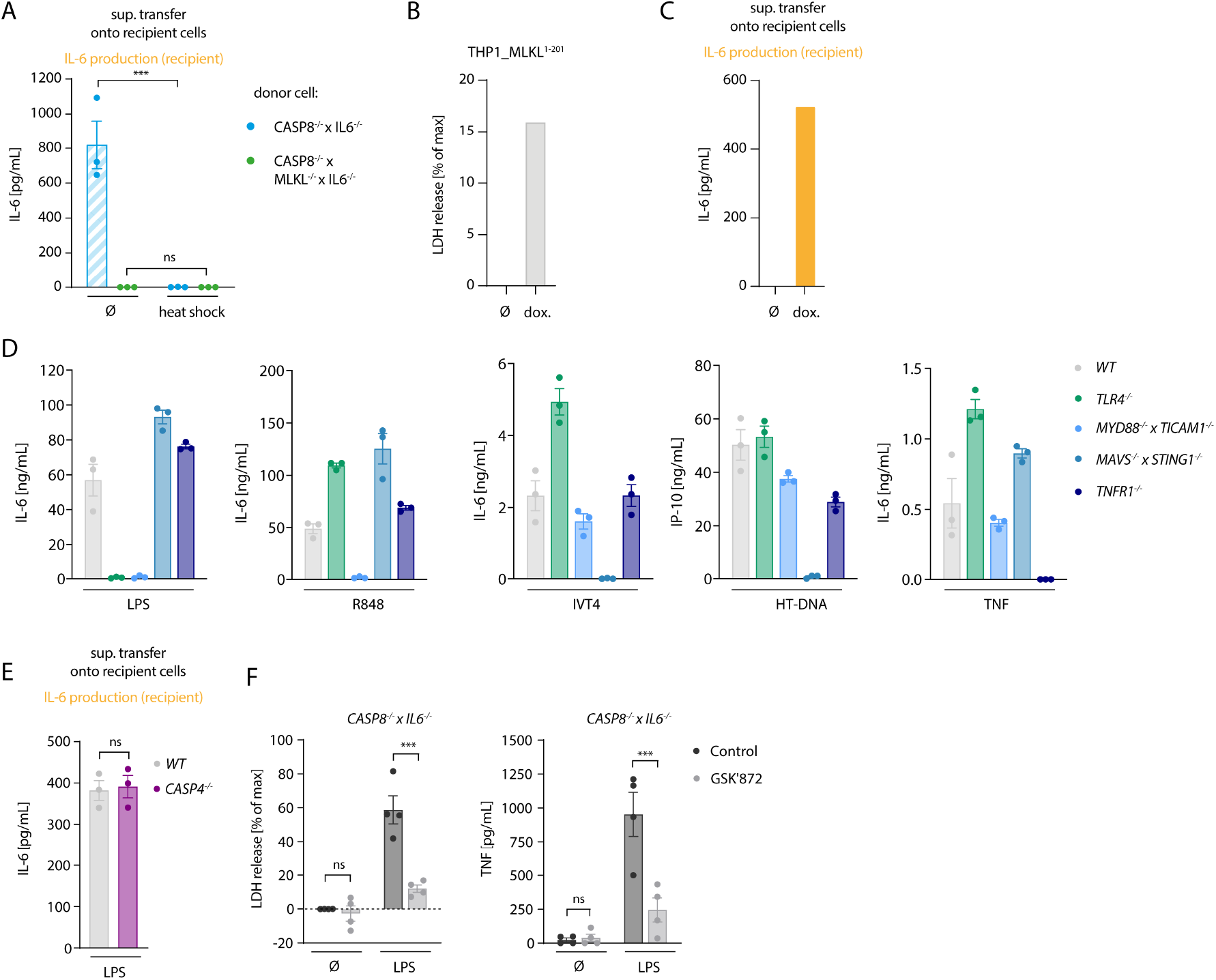
Necroptosis-driven inflammatory responses are mediated by TNF. (A) Supernatant of indicated donor cells (treated for 18 h with 2 ng/ml LPS) was subjected to heat shock for 10 min at 75°C or left untreated and then used to stimulate *TLR4*^-/-^ recipient cells. IL-6 secretion was measured after 24 h. (B) LDH release by THP1 MLKL^1-201^ cells untreated or induced with 1 μg/ml doxycycline for 16 h. (C) IL-6 secretion by *TLR4*^-/-^ BLaER1 cells after stimulation with the supernatant of untreated or doxycycline-induced THP1 MLKL^1-201^ cells. (D) BLaER1 cells of the indicated genotypes were treated with the specified stimuli to phenotypically confirm their deficiency for the respective deleted immune pathways. Either IL-6 or IP-10 secretion was determined. (E) The supernatant of *CASP8*^-/-^ × *IL6*^-/-^ cells stimulated with 2 ng/ml LPS for 18 h was used to stimulate WT and *CASP4*^-/-^ cells for 24 h in presence of 1 μg/ml of the TLR4 inhibitor CLI095. IL-6 release was measured (F) *CASP8*^-/-^ × *IL6*^-/-^ cells were stimulated for 18 h with 2 ng/ml LPS or left untreated in the presence or not of 3 μM the RIPK3 inhibitor GSK’872. LDH and TNF release were measured. Data are presented as one representative experiment out of two (B and C) or as mean values ± SEM of three (A and E) or of four (F) independent experiments or as mean values ± SEM of three biological replicates of one representative experiment out of three (D). Statistics indicates significance by two-way ANOVA (A and F) or an unpaired, two-tailed t-test (E) with a Šidák (A and F) correction for multiple testing: *** P<0.001, ns=not significant. Raw data are available in Figure S2—source data 1.

**Fig. S3.**
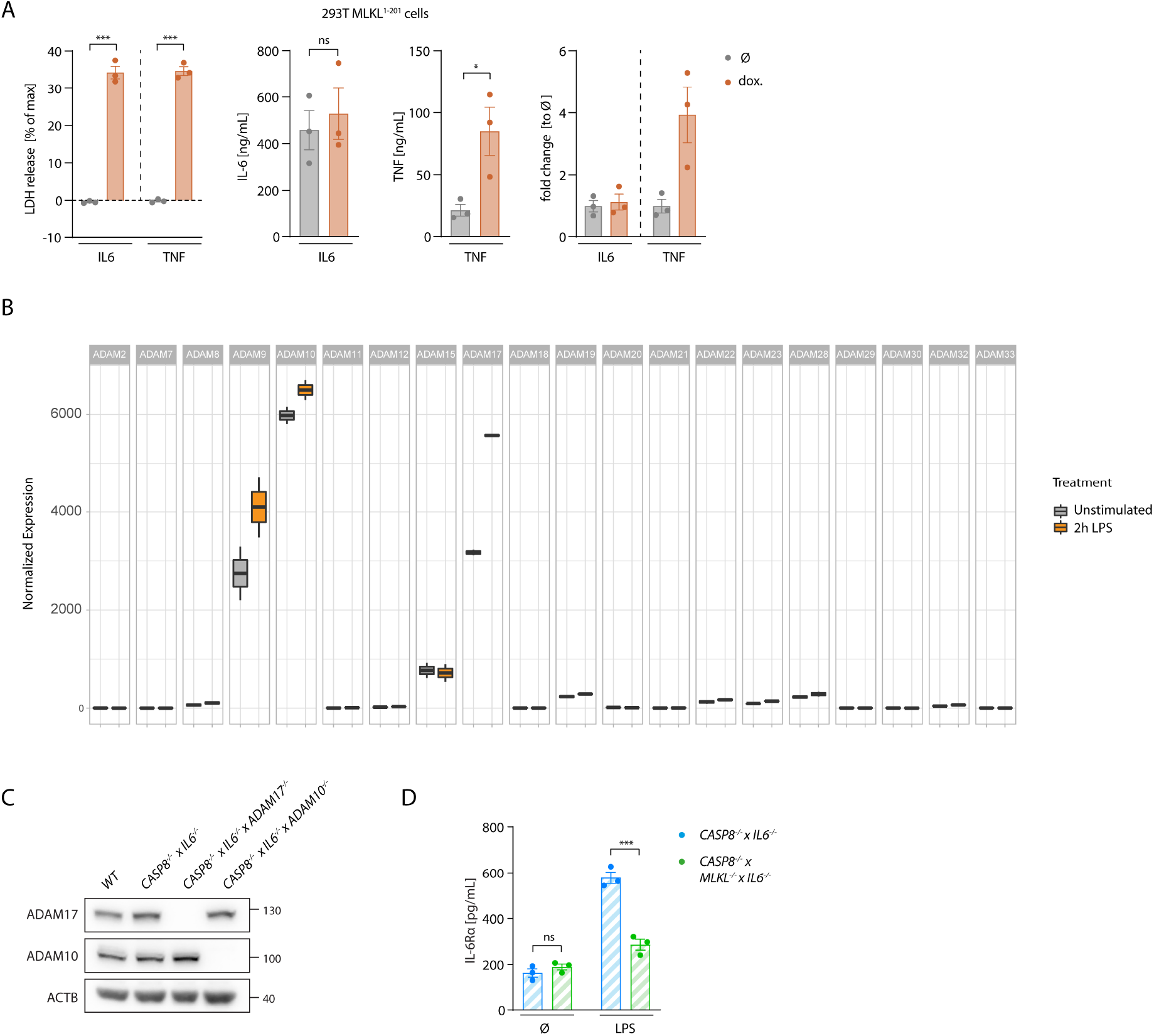
Necroptosis drives ADAM-dependent TNF shedding. (A) 293T cells stably transduced with MLKL^1-201^ were transfected with pRP-TNF or pRP-IL6. The day after cells were treated with 1 μg/ml doxycycline or left untreated for 7 h and LDH and cytokine release were measured. The right panel depicts the cytokine release data as fold change data in relation to the mean value of the data of the untreated cells. (B) Expression of genes of the ADAM family members as from RNA-seq data of WT BLaER1 cells stimulated as indicated. (C) Immunoblotting of ADAM17 and ADAM10 in BLaER1 of the indicated genotypes. (D) IL-6Rα secreted by *CASP8*^-/-^ × *IL6*^-/-^ and *CASP8*^-/-^ × *MLKL*^-/-^ × *IL6*^-/-^ treated with 2 ng/ml LPS for 18 h. Data are depicted as mean ± SEM of 3 independent experiments (A and D). Statistics indicates significance by an unpaired, two-tailed t-test (A) or a two-way ANOVA (D) with a Šidák (D) correction for multiple testing: *** P<0.001, * P<0.05, ns=not significant. Raw data are available in Figure S3—source data 1 and 2.

**Fig. S4.**
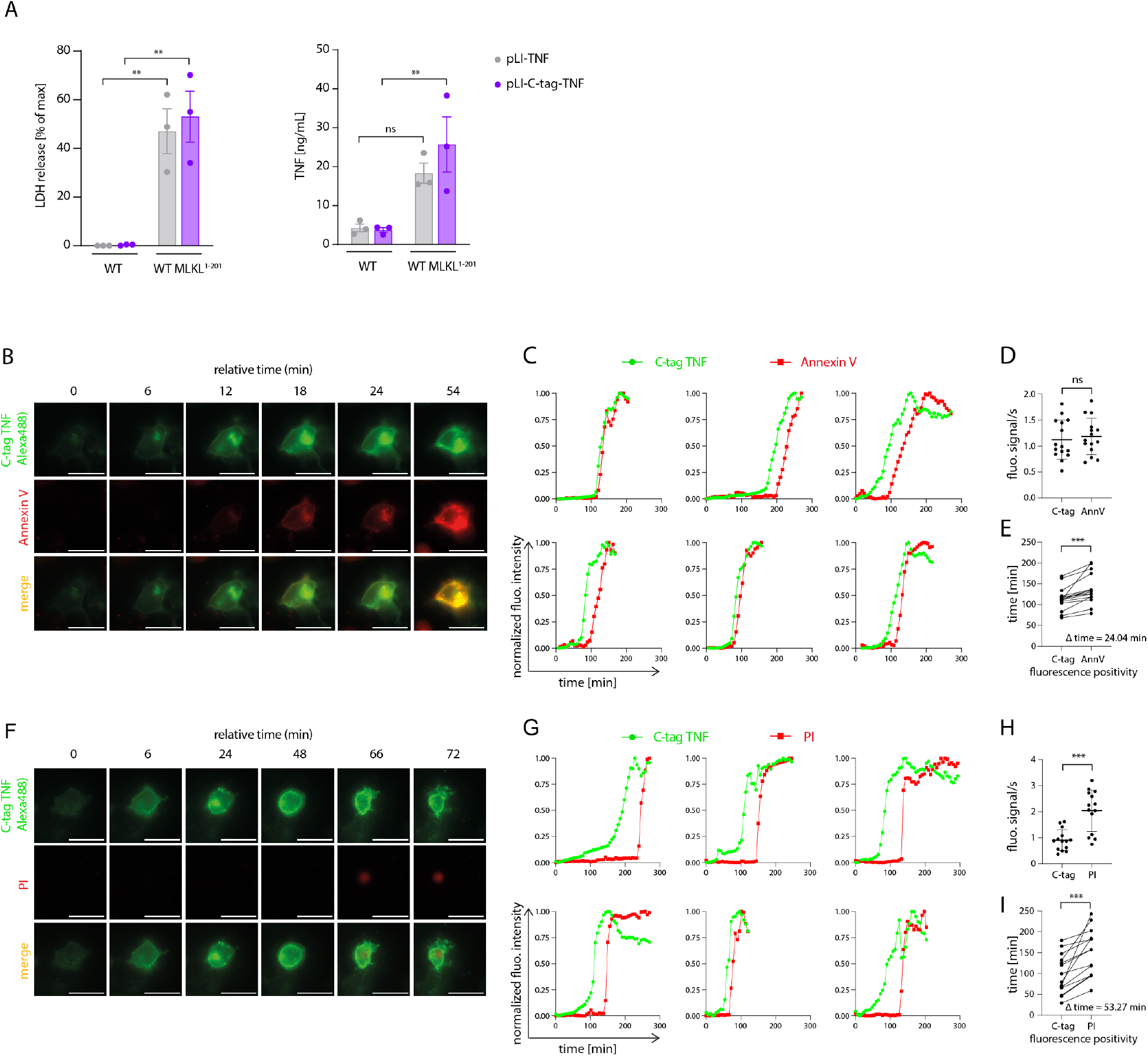
Necroptosis triggers rapid shedding activity. (A) 293T and 293T MLKL^1-201^ cells were transfected with the indicated plasmids. 6 h later, cells were stimulated with 1 μg/ml doxycycline. 18 h later LDH and TNF release were measured. (B-H) 293T MLKL^1-201^ cells were transfected with pLI-C-tag-TNF and induced with 1 μg/ml doxycycline. Imaging of the indicated channels was performed for 4h30 starting 4 h after induction. (B, F) Indicated channels of a representative cell is shown over time. (C, G) Single-cell trajectories of depicted channels over time. (D, H) Calculation of slopes of several cells for the depicted channels. (E, I) Indication of time (in minutes) when cells become positive for the specified fluorescent signal. Representative cells taken from 3 independent experiments were used for the calculations in D, H, E, I. Data are depicted as mean ± SEM of 3 independent experiments (A) or as representative data from three independent experiments (B and F). Statistics indicates significance by a two-way ANOVA (A) or a paired, two-tailed t-test (D, E, H and I) with a Šidák (A) correction for multiple testing: *** P<0.001, ** P<0.01, ns=not significant. Raw data are available in Figure S4—source data 1.

**Fig. S5.**
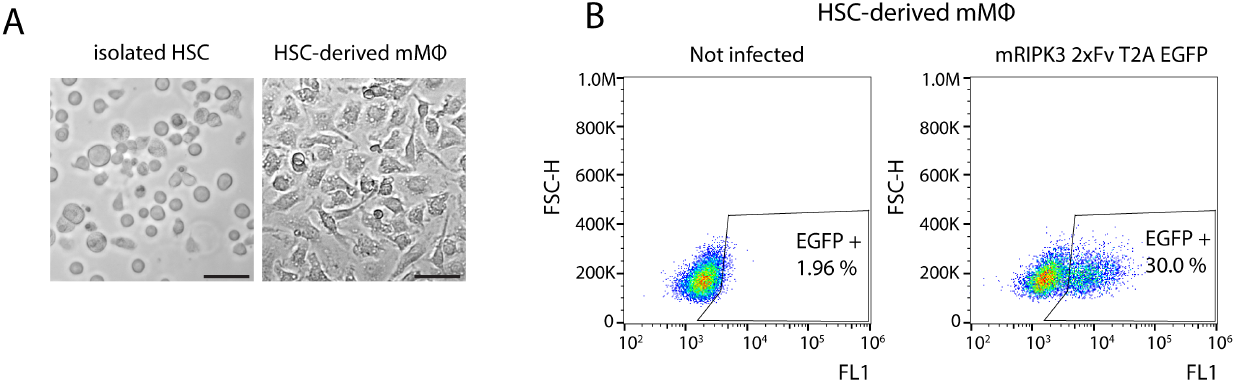
Necroptosis driven TNF release in primary cells. (A) Exemplary image of HSCs on the day of isolation or after differentiation to macrophages. Bar = 25μm. (B) HSCs were transduced with pFUGW-mRIPK3-2xFv-T2A-EGFP the day after isolation or left untransduced. After differentiation, HSC-derived macrophages were analyzed by flow cytometry for the EGFP expression, to assess transduction efficiency.

## Supplementary video 1-3

293T MLKL^1-201^ (video 1), ADAM10^-/-^ × ADAM17^-/-^ MLKL^1-201^ 293T (video 2) and 293T WT (video 3) were transfected with pLI-C-tag-TNF linker mCherry and pEF-BOS-nBFP, stimulated with 1 μg/ml doxycycline and imaged over time (1 image every 6 minutes). Overlay of brightfield, mCherry (TNF) and C-tag TNF is shown.

## Materials and Methods

### Cell culture

BLaER1 ^57^ and THP-1 (DSMZ, ACC 16) cells were cultured in RPMI medium 1640, supplemented with 10% (v/v) FCS, 1% sodium pyruvate (v/v), 100 U/mL Penicillin-Streptomycin (all Gibco). BLaER1 cells were trans-differentiated for 5-6 day in medium containing 10 ng/mL IL-3, 10 ng/mL M-CSF (MPI of Biochemistry, Munich) and 100 nM β-estradiol (Sigma-Aldrich) at a concentration of 7 × 10^4^ cells/well in 96 well plates. Unless otherwise stated, *CASP4*^-/-^ BLaER1 cells were used as genetic background to generate subsequent knock-out clones analyzed in this work. 293T cells (DSMZ, ACC 635) were cultured in DMEM supplemented as above.

### Chemicals, peptides and recombinant proteins

The following reagents were purchased from commercial suppliers: Alexa Fluor® 647 Annexin V (Biolegend, Cat#640911), B/B homodimerizer (Takara Bio, Cat#635058), b-estradiol (Sigma-Aldrich, Cat#E8875-250MG), CLI-095 (InvivoGen, Cat# tlrl-cli95), Collagen R solution 0.2 % (Serva, Cat#47254.02), DNAse I (Thermo Fisher Scientific, Cat#EN0525), Doxycycline hyclate (Sigma-Aldrich, Cat#D9891), eBioscience™ Propidium Iodide Staining Solution (Thermo Fisher Scientific, Cat# 00-6990-50), GeneJuice®Transfection Reagent (Merck Chemicals GmbH, Cat#70967), GeneRuler 1 kb DNA Ladder (Thermo Scientific, Cat# SM0311), GSK’872 (Aobious, Cat#AOB4887), Hepes 1M (Sigma-Aldrich, Cat#H0887-100ML), HT-DNA (Sigma-Aldrich, Cat#D6898), Lipofectamine 2000 (Life Technologies, Cat#11668019), LPS-EB Ultrapure (InvivoGen, Cat#tlrl-3pelps), MCC950 (Sigma-Aldrich, Cat#5381200001), MEM Non-Essential Amino Acids Solution (100×) (Thermo Fisher Scientific, Cat#11140050), PageRuler™ Prestained Protein Ladder (Thermo Fisher Scientific, Cat#26617), Penicillin-Streptomycin (Life Technologies, Cat#15140122), Phorbol 12-myristate 13-acetate (Enzo Life Sciences, Cat#BML-PE160-0005), R848 (InvivoGen, Cat# tlrl-r848), Recombinant Human TNF (Peprotech, Cat#300-01A), RetroNectin® Recombinant Human Fibronectin Fragment (Takara Bio, Cat#T100A), Recombinant Murine Flt3-Ligand (Peprotech, Cat#250-31L), Recombinant Murine IL-3 (Peprotech, Cat#213-13), Recombinant Murine IL-6 (Peprotech, Cat#216-16), Recombinant Murine SCF (Peprotech, Cat#250-03), Recombinant Murine TPO (Peprotech, Cat#315-14), Retronectin (Takara Bio Cat#T100A), Sodium Pyruvate (Thermo Fisher Scientific, Cat#11360088), Streptolysin O (Sigma-Aldrich, Cat# S5265-25KU), Z-VAD-FMK (Peptanova, Cat#3188-v).

### Antibodies

The following antibodies were purchased from commercial suppliers: Alexa-488-conjugated secondary antibody (Thermo Fisher Scientific, Cat# A-11001, RRID:AB_2534069 and Cat# A-11008, RRID:AB_143165), anti-b-actin (C4) HRP (Santa Cruz Biotechnology, Cat# sc-47778 HRP, RRID:AB_271418), anti-Caspase-8 (1C12) (Cell Signaling Technology, Cat# 9746, RRID:AB_2275120), anti-ADAM10, CT (Millipore, Cat# AB19026, RRID:AB_2242320), anti-ADAM17 (D22H4) (Cell Signaling Technology, Cat# 6978, RRID:AB_1082838), anti-IκBα (L35A5) (Cell Signaling Technology, Cat# 4814, RRID:AB_390781), anti-phospho-IkBa (14D4) (Cell Signaling Technology, Cat# 2859, RRID:AB_561111), anti-TNF (D5G9) (Cell Signaling Technology, Cat# 6945, RRID:AB_10859375), CaptureSelect™ Alexa488 anti-C-tag conjugate (Thermo Fisher Scientific, Cat#7213252100), MLKL polyclonal antibody (Thermo Fisher Scientific, Cat# PA5-34733, RRID:AB_2552085), phospho-NF-κB p65 (Ser536) (93H1) rabbit mAb (Cell Signaling Technology, Cat# 3033, RRID:AB_331284), recombinant anti-MLKL (phospho S358) antibody (Abcam, Cat# ab187091, RRID:AB_2619685).

### Derivation of human monocyte-derived macrophages

PBMCs were obtained from healthy volunteers. Informed consent was obtained from all subjects according to the Declaration of Helsinki and approval by the responsible ethical committee (project number 19-238, Ethics committee of the medical faculty of the Ludwig Maximilian University Munich). Monocytes were isolated using the CD14 positive selection method of MACS technology (Miltenyi Biotec, Cat#130-050-201) according to manufacturer’s protocol. Purified monocytes were plated in 96 well plates (0.1 × 10^6^ cells/well) and differentiated into monocytes-derived macrophages (MDMs) for 7-8 days in RPMI medium 1640 supplemented with 10% (v/v) FCS, 1% (v/v) sodium pyruvate, 1% (v/v) non-essential amino acids (Gibco), 100 U/mL Penicillin-Streptomycin and 100 ng/mL M-CSF (Peprotech).

### Plasmids

Cloning of genes of interest into pLI, pRP, pFUGW and pEF-BOS backbones was performed by conventional restriction enzyme cloning. hTNF, MLKL^1-154^ and MLKL^1-201^ were amplified from cDNA of BLaER1 cells. The codon-optimized sequence of hIL6 was ordered from IDT. pMDLg/pRRE was a gift from Didier Trono (Addgene plasmid #12251; http://n2t.net/addgene:12251; RRID:Addgene_12251), pRSV-Rev was a gift from Didier Trono (Addgene plasmid #12253; http://n2t.net/addgene:12253; RRID:Addgene_12253), pLIX_403 (herin referred to as pLI) was a gift from David Root (Addgene plasmid #41395; http://n2t.net/addgene:41395; RRID:Addgene_41395).

### Cell stimulations

Unless otherwise indicated, *CASP8*^-/-^, *CASP8*^-/-^ × *MLKL*^-/-^ and subsequently derived gene-deficient BLaER1 macrophages were stimulated with 2 ng/mL LPS from E.coli (Invivogen) for 18 h. In case of supernatant transfer experiment, the supernatant was then collected, centrifuged at 600 g for 5 minutes and used undiluted to stimulate *TLR4*^-/-^ recipient BLaER1 cells for 24 h. When recipient cells were not in a *TLR4*^-/-^ background, the TLR4 inhibitor CLI095 was added to the collected supernatant at the final concentration of 1 μg/mL. For co-culture experiments, *CASP8*^-/-^ or *CASP8*^-/-^ x *MLKL*^-/-^ BLaER1 macrophages were mixed 1:1 with *TLR4*^-/-^ cells and stimulated for 18 h with 2 ng/mL of LPS. For heath shock experiments, the supernatant of donor cells was heated at 75°C for 10 minutes before stimulation of *TLR4*^-/-^ BLaER1 macrophages. FCS was replaced to a final concentration of 10% after heat shock treatment. The NLRP3 inhibitor MCC950 was used at the final concentration of 5 μM. Stimulation of BLaER1 macrophages with other immune ligands was performed as follows: hTNF (Peprotech) 100 ng/mL, R848 (Invivogen) 1 μg/mL for 16 h. 200 ng/well of IVT4 or Herring testes DNA (HT-DNA, Sigma-Aldrich) were complexed with Lipofectamine 2000 (Life technologies) according to manufacturer’s protocol and cells were stimulated with the mix for 16 h. Primary MDMs were stimulated with 0.2 ng/mL LPS for 8 h. The Caspase-8 inhibitor Z-VAD-FMK (Peptanova) and the RIPK3 inhibitor GSK’872 (Aobious) were added to the cells together with LPS at the final concentration of 20 μM and 3 μM, respectively. For the size exclusion chromatography experiment, THP1 cells previously transduced with pLI-MLKL^1-201^ were differentiated with 100 ng/mL PMA (Sigma-Aldrich) for 24 h before stimulation. Differentiation medium was removed and cells were washed in PBS before incubation in serum-free medium in presence or absence of doxycycline (1 μg/mL) for 16 h. Doxycycline was used to stimulate 293T cells at the final concentration of 1 μg/mL for 7 h on the day after transfection. In case an immunoblot had to be performed, 293T and BLaER1 cells were stimulated respectively in 1% FCS or 3 % FCS containing medium. To induce necroptosis in HSCs-derived macrophages transduced with the dimerizing construct (mRIPK3-2xFv-T2A-EGFP), cells were stimulated with indicated concentrations of B/B homodimerizer (Takara Bio) for 15 h. Streptolysin O was used at the concentration of 8 μg/ml for 7 h to stimulate 293T cells.

### Derivation of macrophages from HSC

Hematopoietic stem cells (HSCs) were obtained from 8-weeks old wild-type C57BL/6 mice. Bone marrow was flushed from femurs and tibias and HSCs were isolated by negative selection with the Direct Lineage Depletion Kit (Miltenyi Biotec, Cat#130-110-470) using MACS technology. Isolated HSCs were cultured in StemPro™-34 SFM (1X) media (Gibco) supplemented with StemPro®-Nutrient Supplement, 100 U/mL penicillin-streptomycin and the following cytokines (all PeproTech): IL-3 (10 ng/mL), FLT3-L (10 ng/mL), SCF (100 ng/mL), TPO (20 ng/mL), IL-6 (20 ng/mL). The day after isolation, HSCs were transduced as described. 24 h later, HSCs were plated in 96 well plate at 0.7 × 10^5^/well and differentiated into macrophages in fully supplemented DMEM with 100 ng/mL M-CSF for 5 days. Medium with fresh M-CSF was added on day 2 or 3 after plating. All mice were handled according to institutional guidelines approved by the animal welfare and use committee of the government of Upper Bavaria.

### CRISPR/Cas9 mediated gene ablation

BLaER1 and 293T cells deficient for specific genes were generated via CRISPR/Cas9 mediated gene ablation as previously described ^58^. Briefly, sgRNAs were designed to target early codons of selected genes. pLK0.1-gRNA-CMV-GFP and a Cas9 expression plasmid (pRZ-BFP-Cas9) were co-electroporated in BLaER1 cells using a Biorad GenePulser device or co-transfected in 293T cells in combination with GeneJuice transfection reagent (Merck). 24 h later, cells positive for respective fluorescent markers were sorted and subjected to limiting dilution cloning. After 3-4 weeks, monoclones were identified and subjected to deep sequencing (Illumina Miseq platform). Clones bearing frameshift mutations were utilized for subsequent experiments. For the generation of TNF Δ 3’ UTR clones, 2 gRNAs targeting the very beginning and the very end of the 3’UTR of the TNF gene were cloned into pMini-U6-gRNA-CMV-BFP-T2A-Cas9 or pMini-U6-gRNA-CMV-mCherry-T2A-Cas9 and co-electroporated in BLaER1 parental cells. Double fluorescent cells were sorted and subjected to limiting dilution cloning. Clones lacking the 3’ UTR were selected by PCR on the target region using the following primers: fwd 5’CCTGGTATGAGCCCATCTATCTG, rev 5’ TTCTTTTCTAAGCAAACTTTATTTCTCGCC (long product = WT, short product =Δ 3’UTR)

### gRNA target sequences

The following genomic regions were targeted to generate knockout cell lines (PAM sequence is depicted in bold letters): ADAM17 GAGCAGAACATGATCCGGA**TGG**, ADAM10 TTTCAACCTACGAATGAAG**AGG**, LTA ACGTTCAGGTGGTGTCATG**GGG**, TNF TGCAGCAGGCAGAAGAGCG**TGG**, MLKL GAGCTCTCGCTGTTACTTC**AGG**, TNFRS1A GCAGTCCGTATCCTGCCCC**GGG**, TICAM1 (TRIF) GGCCCGCTTGTACCACCTGC**TGG**, MAVS ACTTCATTGCGGCACTGAG**GGG**, STING1 GCGGGCCGACCGCATTTGGG**AGG**, MYD88 CTGCAGGAGGTCCCGGCGC**GGG**, CASP4 CTCATCCGAATATGGAGGC**TGG**, IL6 TGTGGGGCGGCTACATCTT**TGG**, TLR4 GATAAAGTTCATAGGGTTC**AGG**, CASP8 GCTCAGGAACTTGAGGG**AGG**, TNF UTR (1) AGGGGGTAATAAAGGGATTG**GGG**, TNF UTR (2) TTACAGACACAACTCCCCTG**GGG.**

### Virus production and cell transduction

For the generation of pseudotyped lentiviral particles, 293T cells were transfected by calcium phosphate co-precipitation with the lentiviral plasmid expressing the desired construct (in a pLI backbone) and the following packaging plasmids: pMDLg/pRRE, pRSV-REV and pCMV-VSVG. 8-14 h after transfection, medium was changed, and cells were incubated for 48-72 h before virus-containing supernatant was harvested and used to transduce target cells. Puromycin selection was applied for 3-4 days before further experimental use. To transduce HSCs with the dimerizing construct (mRIPK3-2xFv-T2A-EGFP expressed in the lentiviral vector pFUGW), 0.5 × 10^6^ cells were plated in a 24-well pre-coated with retronectin (Takara Bio). Virus-containing medium was concentrated by ultracentrifugation and added to cells for 8 h. HSCs were then washed with PBS and transferred to a new 24-well with fresh culture medium. The percentage of transduced cells was evaluated by flow cytometry analysis of EGFP expression on the day of the experimental stimulation.

### Immunoblotting

Cell pellets were lysed in cold RIPA buffer containing a cocktail of protease inhibitors (Roche, Cat# 11697498001). Quantification of proteins was conducted via the Pierce BCA Protein Assay Kit (Thermo Fisher Scientific, Cat#23227). Cell lysates and cell supernatants were denatured in Laemmli Buffer for 10 minutes at 95°C, followed by separation by denaturing SDS-PAGE. Equal amounts of proteins from lysates (15-20 μg) or equal volumes of supernatant were loaded in each lane. In the case of BLaER1 supernatant, proteins were precipitated with methanol/chloroform before denaturation in Laemmli buffer. Proteins were blotted onto a 0.2 μm nitrocellulose membrane, blocked in 5% milk and incubated with indicated primary antibody overnight at 4°C. The day after, membranes were washed and incubated in respective HRP-coupled secondary antibodies for 1 h at room temperature. Upon need, membranes were re-probed with multiple antibodies. Chemiluminescent signal was detected via a CCD camera (Fusion Fx, Vilber).

### Transient transfection of 293T cells

293T cells were transfected with plasmid DNA complexed with GeneJuice following provider’s instructions. For experiments where both immunoblotting and ELISA had to be performed, 0.65 × 10^6^ cells of indicated genotypes were plated in each well of a 6 well plate and transfected the day after, unless otherwise indicated. Co-transfection with 2 plasmids was performed by using 1.5 μg of each plasmid/well. Cells were used for the experiment on the day after transfection. When only ELISA analysis had to be performed, 2 × 10^4^ cells/well were plated in 96-well format and transfected the day after with 200 ng/well of the needed plasmids (100 ng/plasmid in case of co-transfection with 2 plasmids or in case of transfection followed by streptolysin O stimulation). For live cell imaging experiments, 1.7 × 10^4^ 293T cells of the indicated genotypes were plated in collagen R (Serva) coated Ibidi μ-Plate 96 well black (Ibidi) using FluoroBrite DMEM medium (Life Technologies) supplemented as described, with the addition of 10 mM Hepes (Sigma-Aldrich). The day after, cells were transfected with a total of 100 ng DNA/well (50 ng of pLI-C-tag-TNF linker mCherry or pLI-C-tag-TNF + 50 ng pEF-BOS-nBFP, the latter for the expression of a nuclear BFP). 24 h after transfection, cells were stimulated with doxycycline as described.

### Live cell imaging

Imaging was performed starting 4 h after doxycycline treatment to induce the expression of MLKL^1-201^ and/or C-tag TNF/C-tag TNF mCherry. Before imaging, the CaptureSelect™ Alexa Fluor™ 488 anti-C-tag conjugate (Thermo Fisher Scientific) was added to the cell supernatant at a final dilution of 1:480. When necessary, propidium iodide (eBioscience) was added to a final dilution of 1:250 and Annexin V Alexa Fluor™ 647 (Biolegend) was used at 1:125 dilution. Imaging was performed over a period of time of 4h 30 on a Leica DMi8 inverted microscope equipped with a HC PL APO 63x/1.20 W CORR CS2 objective, with a picture taken every 6 minutes. For image analysis, we used *FIJI* (https://imagej.net/) to select regions of interests in individual cells (their motilities were negligible over the observation time) and extract the fluorescence intensity trajectories for the individual fluorescent channels. Different fields of view of at least 3 independent experiments were utilized for the analysis. The mean fluorescence intensities were then plotted as a function of time for single cells and analyzed using *OriginPro2021* (OriginLab). In the case of induction of MLKL^1-201^, an inflection point was visible in the fluorescence intensity trajectories corresponding to the Alexa488 channel (green lines in Fig. 4B). We computed the maximal slopes at the inflection point time as a simple measurement of expression kinetics. In the other cases (WT control and ADAM10^-/-^ × ADAM17^-/-^ MLKL^1-201^ cells), we calculated an average slope by performing a linear regression of the whole fluorescence intensity trajectories.

### Size exclusion chromatography (SEC)

THP1 cells were stimulated as described above in serum-free medium. Supernatant from doxycycline-induced or uninduced cells (40mL per condition) was collected and concentrated to 500 μL using Amicon Ultra-15 filters with 10 kDa cutoff (Merck Millipore). The concentrated samples were loaded on a Superose 6 10/300GL column (GE Healthcare) and an isocratic run (1.2 column volume) was conducted on an ÄKTA Basic instrument with a flowrate of 0.3 mL/min. Degassed and filtered PBS was used as running buffer. 500 μL fractions were collected and utilized for 14 h stimulation of TLR4^-/-^ BLaER1 recipient cells. Upon stimulation, fractions were diluted 1:1 with fully supplemented RPMI medium, adjusting the FCS content to 10% v/v.

### RNA sequencing

Differentiated BLaER1 cells were left untreated or LPS stimulated for 2 h. Next, 3×10^6^ cells were lysed in 1ml Trizol and library preparation performed as described elsewhere ^59^. The libraries were sequenced on an Illumina HiSeq1500 device. Reads were aligned to the human reference genome GRCh38 using STAR v.2.5.1 ^60^ and counted using the GenomicAlignments R package ^61^. Post mapping analysis was conducted in R ^62^. Normalization and scaling were performed using DESeq2’s ^63^. GGplot2 ^64^ was used to visualize results.

### RT-qPCR

RT-qPCR analysis of LPS stimulated BLaER1 cells of indicated genotypes was performed as follows. RNA from BLaER1 cells was purified via the Total RNA purification mini spin kit (Genaxxon, Cat#S5304.0250) and DNA digestion was performed with DNAse I (Thermo Fisher Scientific) incubation for 30 minutes. 700 ng of RNA was used for cDNA synthesis by RevertAid Reverse transcriptase (Thermo Fisher Scientific, Cat#EP0442) with oligo dT primers. qPCR reaction was performed with the PowerUp™ SYBR™ Green Master Mix (Thermo Fisher Scientific) in presence of primers specific for genes of interest.

### ELISA

hIL-6 (BD Biosciences, Cat#555220), hTNF (BD Biosciences, Cat#555212), hIL-8 ELISA (BD Biosciences, Cat#555244), hIP-10 (BD Biosciences, Cat#550926), mmTNF (BD Biosciences, Cat#558534) and hIL-6Rα ELISA (R&D Systems, Cat#DY227) were performed according to provider’s protocol.

### LDH

Pierce LDH Cytotoxicity Assay Kit (Thermo Fisher Scientific, Cat#C20301) was performed according to manufacturer’s instructions. Relative LDH release was calculated as LDH release [%] = 100 * (measurement – unstimulated control)/ (lysis control – unstimulated control).

### Quantification and statistical analysis

Statistical significance was determined as explained in the figure legends with the respective post hoc corrections for multiple testing, if relevant. If multiple comparisons are depicted with one comparison bar, the major tick of the comparison bar indicates the reference data to which the statements regarding the level of significance are made. Statistical analysis was performed with GraphPad Prism 8. *** p < 0.001, ** p < 0.01, * p < 0.05, ns = not significant.

### Data and materials availability

BLaER1 RNA-Seq data can be accessed at Sequence Read Archive with the following accession number: PRJNA716479. All data including Source Data for Figs. 1 - 7 and Fig. S1 - S4 are provided with the paper Further information and requests for resources and reagents should be directed to and will be replied to by the corresponding authors.

## Acknowledgments

We kindly acknowledge Andreas Wegerer and Larissa Hansbauer (Gene Center, LMU) for great technical support, Niklas Schmacke (Gene Center, LMU) for deep sequencing, Joshua Kie from the BioSysM FACS Core Facility (Gene Center, LMU) for cell sorting. This work was funded by the Deutsche Forschungsgemeinschaft (DFG, German Research Foundation) CRC 1403 (project number 414786233) to VH.

## Author contributions

Conceptualization, F.P., M.M.G. and V.H.; Formal analysis, G.K. and C.J.; Investigation, F.P., D.N.; Resources, M.M.G.; Writing, F.P. and V.H. with input from all authors; Funding acquisition, V.H.; Supervision, V.H.

